# Plasmon-Coupled Photocapacitor Neuromodulators

**DOI:** 10.1101/2020.02.18.953604

**Authors:** Rustamzhon Melikov, Shashi Bhushan Srivastava, Onuralp Karatum, Itir Bakis Dogru, Houman Bahmani Jalali, Sadra Sadeghi, Ugur Meric Dikbas, Burak Ulgut, Ibrahim Halil Kavakli, Sedat Nizamoglu

## Abstract

Efficient transduction of optical energy to bioelectrical stimuli is an important goal for effective communication with biological systems. For that plasmonics has significant potential via boosting the light-matter interactions. However, plasmonics has been primarily used for heat-induced cell stimulation due to membrane capacitance change (i.e., optocapacitance). Instead, here we demonstrate that plasmonic coupling to photocapacitor biointerfaces improves safe and efficacious neuromodulating displacement charges for an average of 185% in the entire visible spectrum while maintaining the Faradaic currents below 1%. Hot-electron injection dominantly leads the enhancement of displacement current at blue spectral window, and nanoantenna effect is mainly responsible for the improvement at red-spectral region. The plasmonic photocapacitor facilitates wireless modulation of single cells at 3-orders of magnitude below the maximum retinal intensity levels corresponding to one of the most sensitive optoelectronic neural interfaces. This study introduces a new way of using plasmonics for safe and effective photostimulation of neurons and paves the way toward ultra-sensitive plasmon-assisted neurostimulation devices.

## Introduction

Extracellular stimulation is the basis for the communication with the biological systems to recover lost functionalities [1], understand cellular processes [2] and switch neural networks [3]. For that light offers a non-invasive neuromodulation trigger with high spatial and temporal precision [4–8]. Optogenetics introduces light-sensitive opsins into the membrane via a viral transfection, but genetic modification currently raises concerns on their clinical use [9, 10]. Alternatively, geneless optical stimulation of cells has high promise to control the neural activity [11, 12]. For that optoelectronic biointerfaces can generate photo-excited electrons and holes that can be manipulated for Faradaic, capacitive and thermal photostimulation of neurons.

Plasmonics has high potential to increase the performance of optoelectronic biointerfaces. In principle, plasmonic nanostructures can concentrate the incoming radiation to a subwavelength spatial profile due to localized surface plasmon resonance (LSPR) [13], and may boost the light-matter interactions for sensitive transduction of optical signal to bioelectrical stimuli. In addition, light-induced plasmon energy can be transferred to the conduction band of the nanostructure that can produce energetic electrons, known as hot electrons, and these charges may be controlled for photostimulation of neurons. However, despite its high potential for extracellular stimulation, plasmonics has been mainly used for photothermal transmembrane modulation of neurons [14–18]. Light is converted to heat energy due to decay of plasmon oscillations, and the resultant temperature variation induces a membrane potential change through a transient membrane capacitance shift [14]. Recently, some evidence showing that metal decoration can support increase on capacitive and Faradaic currents were also reported [8]. In this study, we demonstrate that the coupling of plasmons to photocapacitors can improve the charge injection of the neuromodulating displacement currents in the entire visible range. We proved that this broadband enhancement ability mainly stems from hot-electron injection at blue and local field enhancement at red spectral regions. Finally, the plasmon-assisted capacitive photocurrent modulates transmembrane potential at a single-cell level.

## Results

### Structure of plasmonic biointerface

Capacitive charge-injection, which is a safe method for stimulation of neurons [8], is based on the electromagnetic attraction and repulsion of the ions in the biological medium due to charge movement in the stimulating electrode. For that the fabrication of plasmonic biointerface architecture (Fig. 1a) started with the sequential deposition of the ZnO and bulk heterojunction composite PTB7-Th:PC71BM layers on ITO (indium tin oxide) substrate. According to the cross-sectional scanning-electron microscopy (SEM) (Fig. 1a, left), the thickness of the ZnO and PTB7-Th:PC71BM layers correspond to 52 nm and 104 nm, respectively. Afterward, gold (Au) thin film is deposited and annealed to form self-assembled nanoislands [19, 20]. The surface morphology of plasmonic biointerface confirms the formation of gold nanoislands (AuNIs) on the photoactive layer (Fig. 1a, right) which leads to a plasmonic peak around 626 nm (Fig. 1b inset). The broad plasmonic band stem from low annealing temperature [19] which is required to retain optical properties of organic photoactive layer. The absorption band of the plasmonic biointerface covers all the visible range, which is useful for the conversion of light within the entire visible range to bioelectrical stimuli (Fig. 1b).

**Figure 1.**
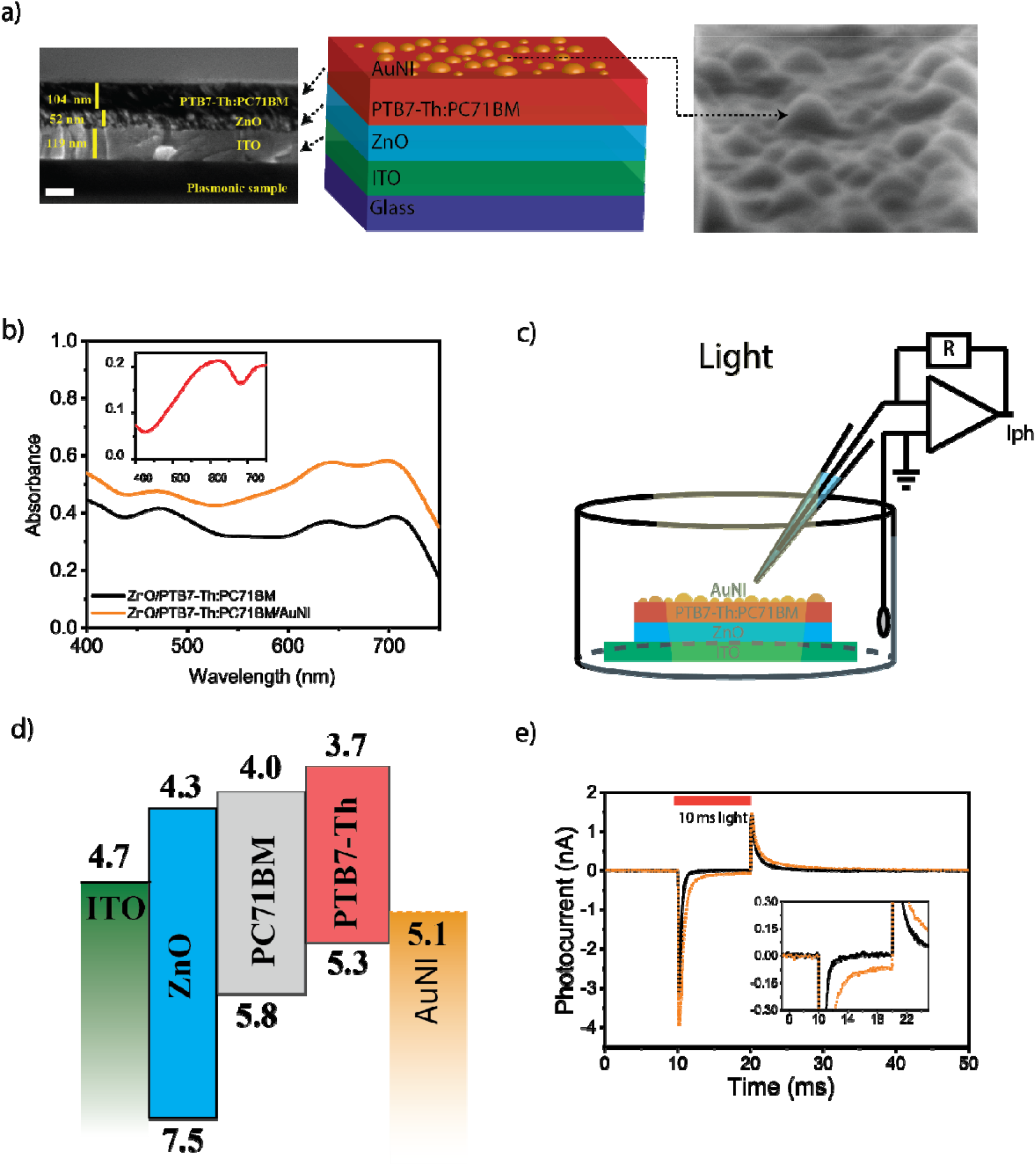
(a) (Left) Cross-sectional scanning electron microscopy (SEM) image of the plasmonic biointerface (scale bar, 100 nm). (Middle) Schematic device structure of the plasmonic (ITO/ZnO/PTB7-Th:PC71BM/AuNI) biointerface. (Right) SEM image of nanoislands on the plasmonic biointerface. (b) Absorption spectrum of ZnO/PTB7-Th:PC71BM (black line) and ZnO/PTB7-Th:PC71BM/AuNI (orange line) thin films. The inset shows absorbance of AuNI layer (red line), which is calculated by extracting it from the control biointerface without gold layer that has a smooth surface due to the absence of nanoislands (Fig. S1a and S1b). (c) Schematic of the photocurrent measurement set-up. The patch pipette is kept close to the surface and the current is measured with a voltage-clamp mode. (d) Energy band diagram of the plasmonic biointerface (with respect to vacuum). (e) Photocurrent of control (black line) and plasmonic (orange dotted line) biointerfaces illuminated under green light at 8.8±0.2 × 10^16^ photons s^−1^.cm^−2^ with 10 ms pulse-widths. The inset zooms on low photocurrent levels. The capacitive current amplitude is defined as the maximal current amplitude reached after the light onset. Faradaic current is defined as the current amplitude for 9LJms after the start of illumination (Fig. S5).

### Photocurrent analysis

We investigate the photocurrent response of the plasmonic biointerface in aCSF (artificial cerebrospinal fluid) solution. We use a patch-clamp system (HEKA, EPC800) in voltage clamp mode (with ~ 6 MΩ patch pipette tips) under the free-standing condition (Fig. 1c). Since the optoelectronic processes are quantal, the biointerfaces were pumped under illumination of blue, green and red lights by keeping the photon counts at the same level (Fig. S2-S3), which correspond to 8.8±0.2 × 10^16^ photons s^−1^.cm^−2^ with 10 ms pulse-widths. The plasmonic biointerface generates several nanoamperes of photocurrent in aCSF in the entire visible spectrum (Fig. S4). Among different excitation wavelengths, green spectral window shows the highest photocurrent for both biointerfaces due to higher absorption strength and external quantum efficiency of the photoactive blend [21] in comparison with the blue and red excitation (Fig. S4). Two spikes were observed at the onset and offset of light illumination. These two spikes (i.e., onset and offset peaks) show that the charge-transfer transfer mechanism is based on the capacitive charge-transfer [22].

Here the current generation mechanism is controlled by the energy band alignment of the layers in the biointerface (Fig. 1d). Photogenerated charge carriers initially dissociate in the photoactive blend and electron transport toward ZnO layer takes place. Holes are mainly localized in the proximity of the electrolyte interface that induces a displacement current. Hence, the initial hole accumulation near electrolyte interface generates onset capacitive photocurrent from the biointerface to the electrolyte (Fig. 1e). After the light is turned off, the decrease of the hole concentration due to recombination leads to another offset capacitive spike in the opposite direction.

We compared the light-induced current and charge generation characteristics of the biointerface with the control biointerface. For the analysis we identified Faradaic and capacitive contributions at blue, green and red spectral windows under the same photon counts (Fig S5. Eqn. S1-S4). The total capacitive charge transfer to the electrolyte is improved for 185.6%, 163.2% and 207.9% at blue, green and red spectral windows, respectively, which correspond to an average increase of 185% in the entire visible spectrum (Fig. 2a, Table S1, S2). Furthermore, the plasmonic biointerface enhanced the peak photocurrents at the level of 34.5%, 28.0% and 25.7% for blue, green and red illumination, respectively (Fig. 2b) (Table S3, S4). We observed that since LUMO energy of PTB7-Th is above water oxidation energy, the Faradaic current due to hole transfer to electrolyte is significantly suppressed that correspond less than 1% of the total photocurrent (Fig. S6). The linear response of the biointerface to the light intensity indicates a single-photon-absorption induced charge carrier generation (Fig. 2c, Fig. S7). We also analyzed the possible contribution by the photothermal effects and since the intensity levels that the plasmonic biointerface operates several orders of magnitude lower than the heat-induced neurostimulation, we did not observe any temperature variation in the solution even under 170 mW.cm^−2^ (Fig. S8). Furthermore, we varied the pulse width between 50 μs-100 ms, and we observed that the peak capacitive currents are well maintained for the pulses longer than 200 μs, which are sufficiently short to elicit targeted electrophysiological processes. The decrease of the photocurrent amplitude below 200 μs stems from the strong overlap between the onset and offset peaks of the capacitive spikes (Fig. 2d, Fig. S6).

**Figure 2.**
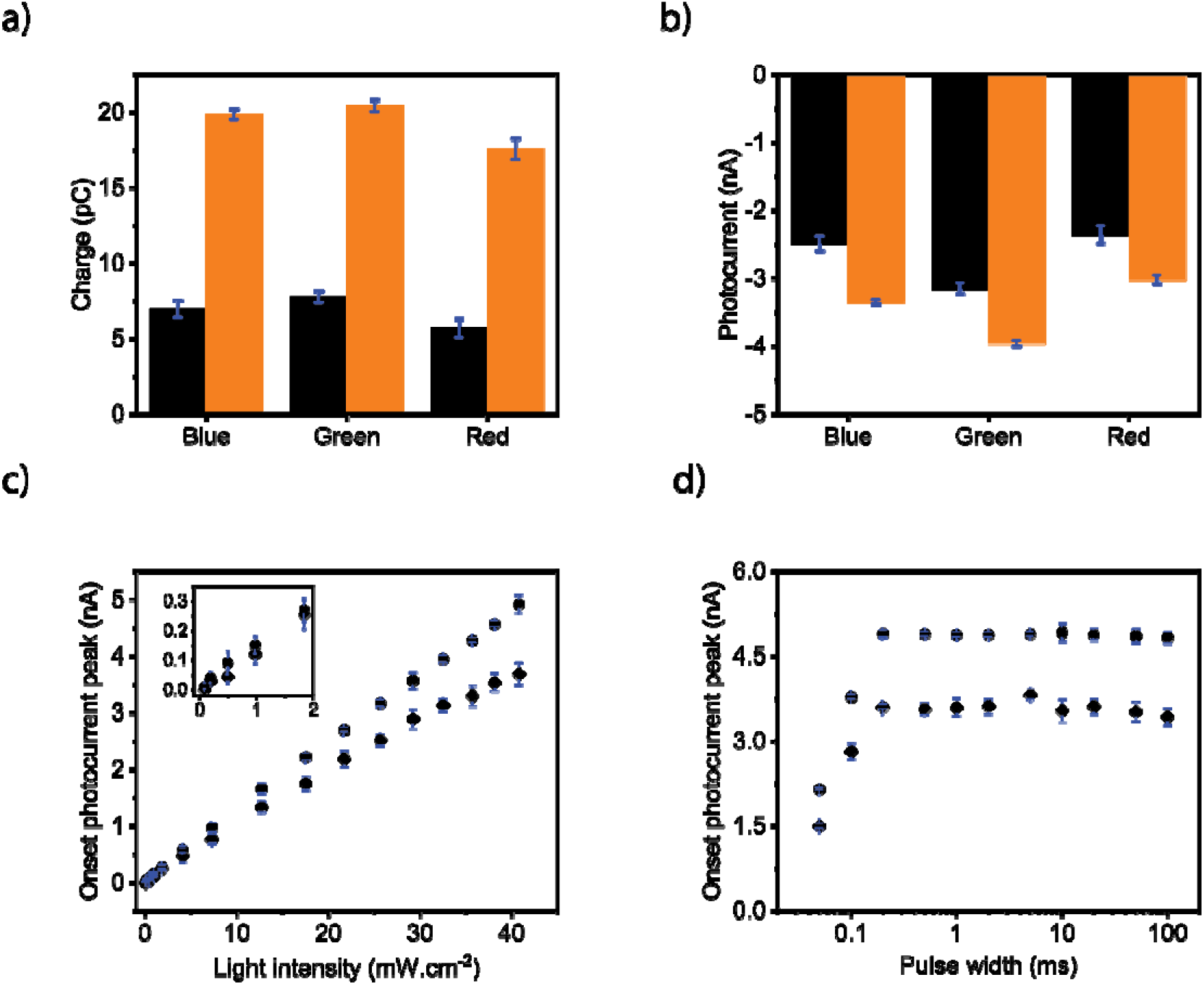
(a) Injected amount of charges and (b) current peaks of the control (black) and plasmonic biointerfaces (orange) at 8.8±0.2 × 10^16^ photons s^−1^.cm^−2^ with 10 ms pulse-widths. (c) Magnitude of peak of capacitive photocurrentcurrents of the control (rhombus dots) and plasmonic (circle dots) biointerfaces under different light intensity levels. (d) Magnitude of the peak capacitive currents generated by the control (rhombus dots) and plasmonic (circle dots) biointerfaces under different pulse widths having light intensity at 8.8±0.2 × 10^16^ photons s^−1^.cm^−2^.

### Electrochemical investigation of plasmonic enhancement

We investigate the enhancement of the displacement current by the plasmonic biointerface using electrochemical impedance spectroscopy. For that we did 3-probe measurements in aCSF media using Ag/AgCl as a reference electrode, platinum as a counter electrode and biointerface as the working electrode [23]. The physical mechanisms at the various interfaces are probed in response to a small AC perturbation of 10 mV (V_rms_) varied over a frequency range from 1 Hz to 0.1 MHz at zero DC bias. Impedance response of the control and plasmonic biointerfaces are semicircle in the high to mid frequency region and depressed semicircle with a linear extension in the low frequency region in the complex plane (Fig3 (a)-(c)). The Bode plot at high frequency is generally interpreted as the electrolyte region and the kHz frequency region corresponds to the double layer formed at the photoelectrode/electrolyte interface (Fig. S9). Bode phase angle (◻ 56°) is found similar for both biointerfaces under blue and green light illumination, and it is interestingly observed higher (◻ 62°) for the plasmonic biointerface under red light illumination (Fig 3(d), (e)).

**Figure 3.**
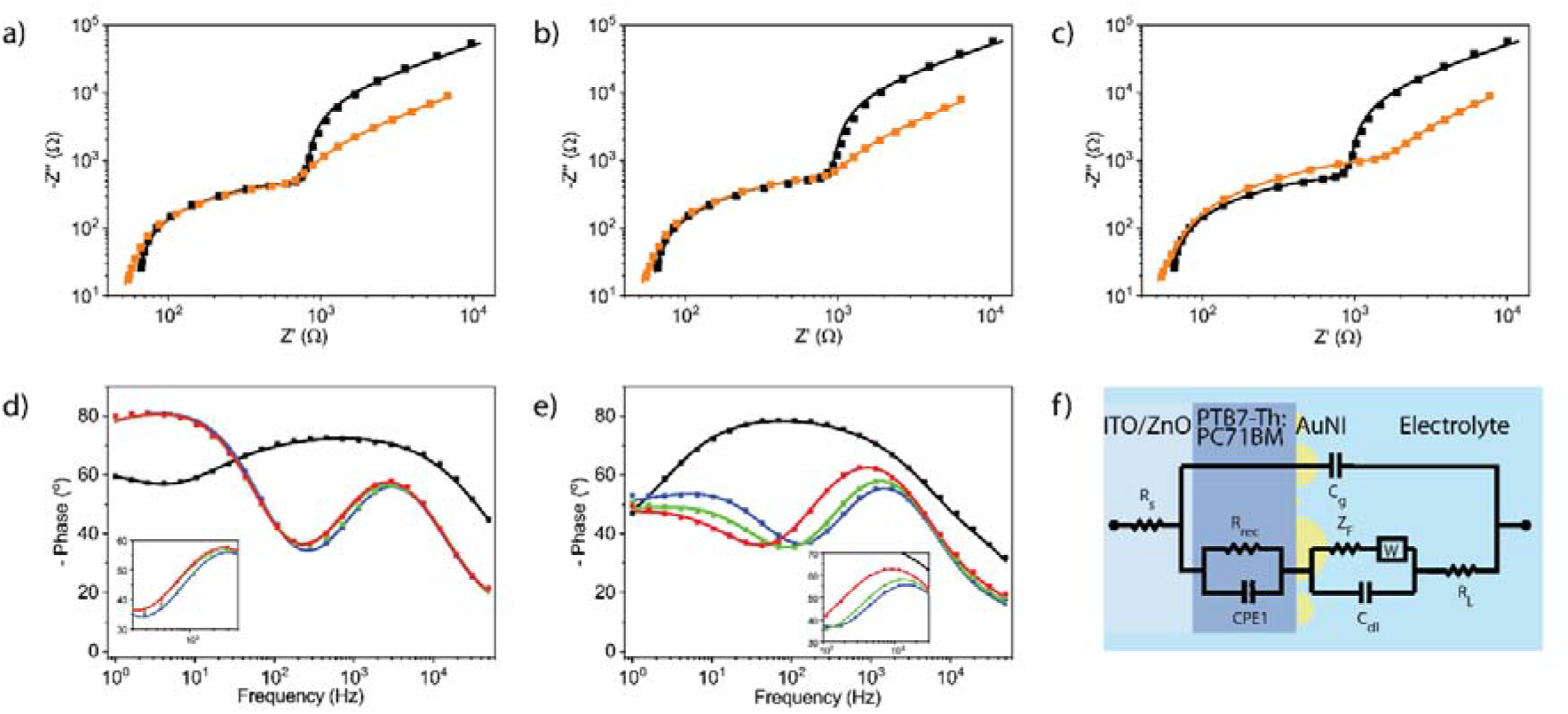
Electrochemical characterization of the control and plasmonic biointerfaces. Nyquist plot of the control (black) and plasmonic biointerfaces (orange) (a) under blue, (b) green and (c) red excitation. Bode phase of (d) the control and (e) plasmonic biointerface. The inset zooms between 50-5000 Hz. (f) Circuit diagram fitting the Nyquist and Bode plot for both the control and plasmonic biointerfaces.

To correlate with the physical mechanisms in the plasmonic biointerface we analyzed the frequency response and fitted the equivalent circuit model (χ^2^=0.01-0.03) comprised of metal-insulator-metal structure (Fig. 3f). C_g_ represents the geometrical capacitance of the device and R_s_ is assigned to the electrical resistance that present at metal electrodes and electrode/contact layer/active layer interfaces. The recombination resistance (R_rec_) in parallel with constant phase element (CPE1) corresponds to the charge dynamics in the photoactive bulk heterojunction. The random characteristics of the blend active layer in organic bulk heterojunction can be attributed to the distribution of relaxation times and are related to the impedance as 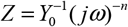, where Y_0_ is the CPE coefficient and the ideality factor n − 0 for pure resistive and 1 for pure capacitive behavior – is the characteristic of the distribution of relaxation times [24]. The plasmonic photoelectrode/electrolyte interface is defined by the parallel combination of Warburg element (W) with double layer capacitance (C_dl_). Using the fitted macro-electrical parameters (Table S5), the optoelectronic parameters of the biointerfaces such as effective lifetime (τ_*n*_) and mobility (*μ*_*n*_) of charge carriers are determined (Table S6).

We observed that while the recombination resistance (R_rec_) is similar both for control and plasmonic biointerfaces under blue and green light, it increases approximately 2-times (R_rec_ =1.6 kΩ) for the plasmonic biointerface under red light in comparison with the control biointerface. This leads to a rise of effective recombination lifetime and indicates a higher number of photo-activated charge generation. Since the peak plasmonic absorption by the gold nanoislands is at red-spectral range, the presence of gold nanoislands at the biointerface can increase the electron concentration by near-field enhancement of the incoming electromagnetic wave, which is observed by the corresponding recombination lifetime increase (Table S6).

In blue spectral region, while the recombination resistance (R_rec_ = 0.66 kΩ) and effective lifetime (τ_n_ = 0.3 ms) remain at similar level with the control biointerface, there is an increase in electron mobility for the blue excitation compared to the red excitation of the plasmonic biointerface. This means that electron injection becomes more probable in blue spectral range. According to the band energy diagram, the energy of blue photons can lead to the excitation of the electrons to an energy level, which is higher than the barrier between gold nanoislands and HOMO level of donor the molecules (PTB7-Th). Hence, the hot electrons with high kinetic energy have higher probability to reach the ITO/ZnO interface [25]. This shows that while at red spectral region nanoantenna effect is dominantly responsible for near-field enhancement, at blue spectral range hot carrier injection becomes the leading factor. Furthermore, the increase of the double layer capacitance at all wavelengths from 2.1 to 3.1 μF/cm^2^ supports the plasmonic enhancement of capacitive photocurrent at each color.

### Biocompatibility

To test the biological and electrophysiological activity we used SHSY-5Y cells. These kinds of non-spiking cells such as N2A, oocytes are already used to prove the neuromodulation ability [26, 27]. Initially, we tested the mitochondrial activity with 3-(4,5-dimethylthiazol-2-yl)-2,5-diphenyltetrazolium bromide (MTT) and lactate dehydrogenase (LDH) assays to determine the toxicity of the plasmonic biointerfaces on SHSY-5Y cell line. In these assays we used ITO substrate as the control group. Plasmonic biointerfaces did not exhibit any toxicity on the cells by either assay (Fig. 4a, 4b). We also evaluated the effect of plasmonic biointerfaces on cell growth by investigating morphology of the SH-SY5Y cells via fluorescence microscopy. Cells grown on the samples were fixed with 4% paraformaldehyde (PFA). Nucleus of the cells were stained with 4’,6-diamidino-2-phenylindole (DAPI) and the cytoplasm was visualized by anti-beta-III tubulin immunolabeling. As can be seen in Figure 4c, cells grown on biointerfaces exhibited comparable morphology with the cells grown on ITO control. All these results suggested that plasmonic biointerfaces are suitable for biological experiments with SHSY-5Y cells.

**Figure 4.**
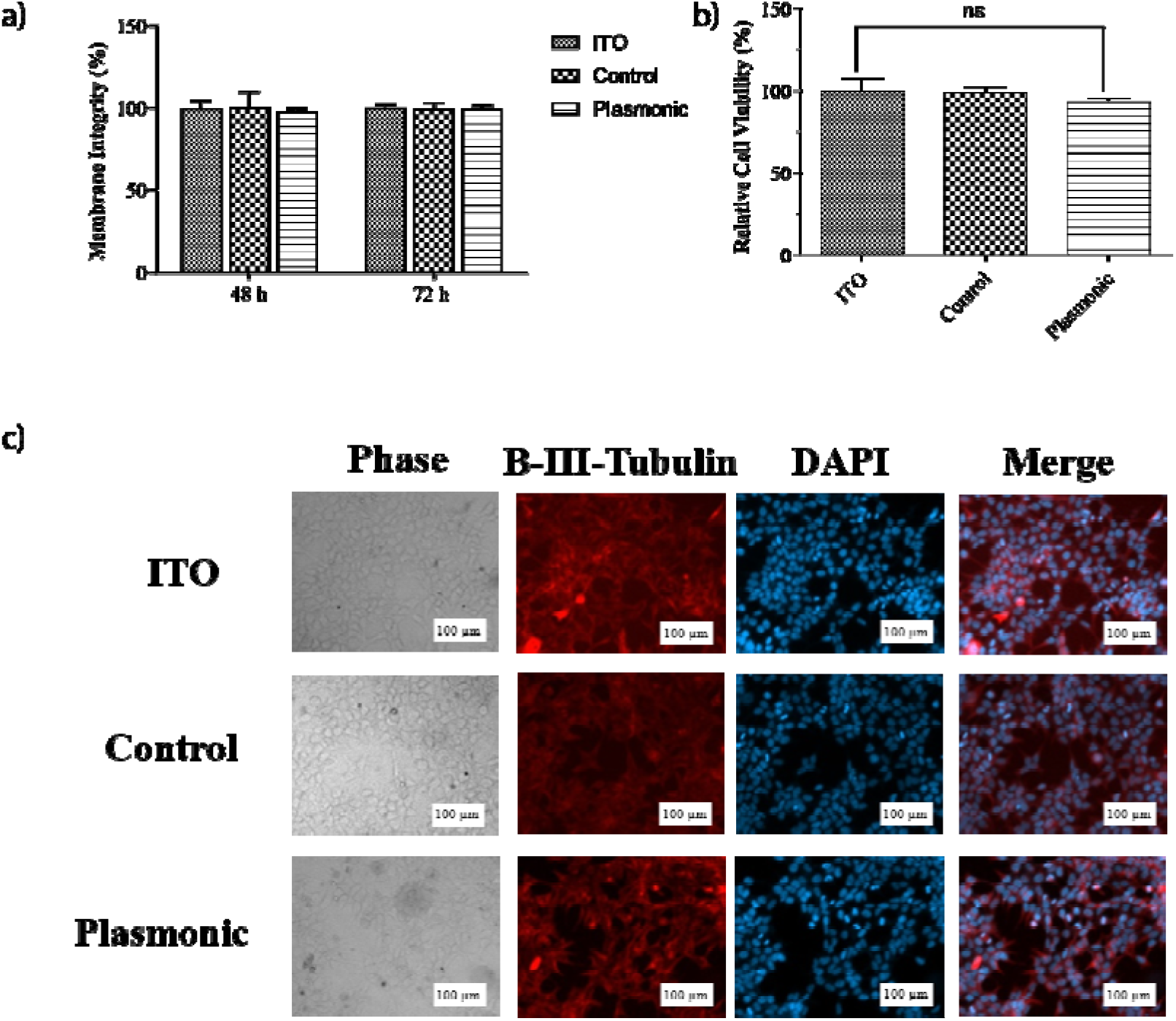
(a) MTT assay, assessment of the effect of plasmonic biointerfaces on mitochondrial activity of SHSY-5Y cells. Cell viability on biointerfaces was presented relative to ITO control. Results are presented in a column graph plotting the mean with standard error of the mean (SEM). Experiments were performed with at least three technical replicates and repeated three times (n=3). An unpaired two-tailed t test was performed to determine the level of significance. *p < 0.05 was considered as statistically significant, nonsignificant differences are presented as ‘ns’. (b) LDH leakage assay, assessment of membrane integrity of the cells grown on biointerfaces. Experiments were performed with at least three technical replicates and repeated three times (n=3). (c) Immunofluorescence imaging, effect of biointerfaces on the morphology of SHSY-5Y cells. Morphology of the cells grown on biointerfaces was visualized by fluorescence microscope after beta-III tubulin immunolabeling and DAPI staining (scale bar: 100 μm).

### Photostimulation of neurons

To investigate the membrane potential variation, we perform patch clamping in whole-cell configuration (Fig. 5a). The I-V characteristics of SHSY-5Y cells grown on both control and plasmonic biointerfaces under dark condition shows that the SHSY-5Y cells exhibit typical resting membrane potential in the range −20 mV and −40 mV (Fig. 5b). The SHSY-5Y cells exhibit quasi-linear response around the resting membrane potential on both biointerfaces. Under 10-ms green light (1 mW.cm^−2^ −40 mW.cm^−2^) an initial depolarization of a single-cell membrane depolarization is followed by a hyperpolarization (Fig. 5c). Since the current direction of the first capacitive peak current is from the biointerface toward the substrate in the attached membrane part, the current leads to a depolarization in the free membrane. In contrast, the second peak of the capacitive current leads to a hyperpolarization of the free membrane (see Fig. S10). Similar membrane behavior under illumination is also observed for green and red illumination as well (Fig. S11). Moreover, under repetitive excitation the peak membrane potential change is also stable under different illumination colors (Fig. S12, Fig. 6e and Table S8).

**Figure 5.**
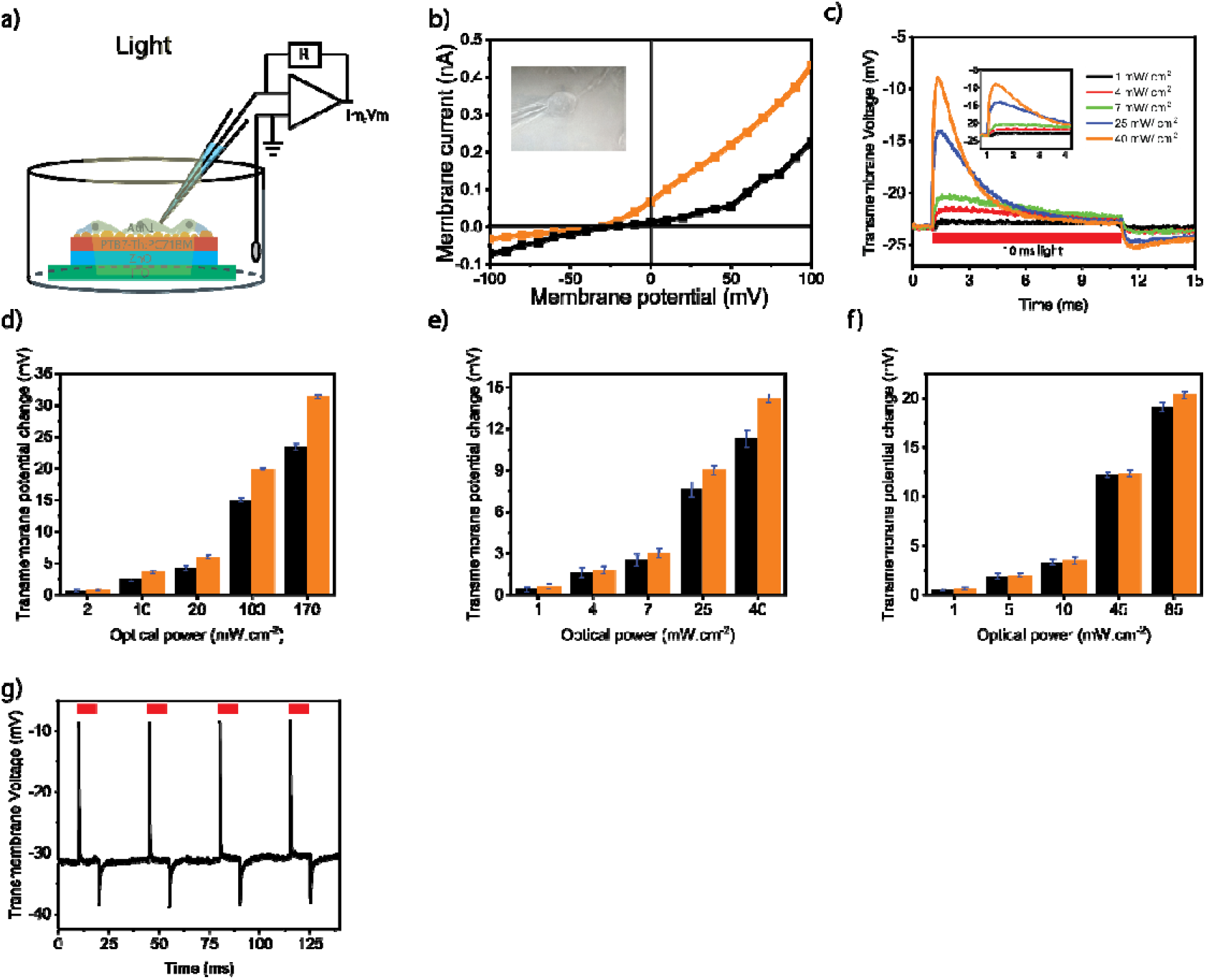
(a) Schematic of the whole-cell patch clamp recording configuration of the biointerface in wireless and free-standing mode. The cells are grown on the biointerfaces and adhere on the surface. (b) Current-voltage characteristics of a typical SHSY-5Y cell adhered on the control (black line) and plasmonic (orange line) biointerfaces. The inset shows photograph of a SHSY-5Y cell on the plasmonic biointerface patched with a glass micropipette. (c) Transmembrane potential variation of SHSY-5Y cells on the plasmonic biointerface under green light illumination with 10 ms pulse width. (d) – (f) Transmembrane potential variation of SHSY-5Y cell on the control (black) and plasmonic (control) biointerfaces under blue (d), green (e) and red (f) light illumination with 10 ms pulse width. (g) Transmembrane potential under 10-ms pulse train.

In comparison with the control biointerface, we observed a 37.3 % enhancement in the peak depolarization of the transmembrane potential of the SHSY-5Y cells for blue excitation (Fig. 5d). While the excitation is shifted to green and red colors, we observed lower peak enhancement of 25.3 % and 7.7 %, respectively (Fig 5(e)-5(f), Eqn. S5-S6 and Table S7). This shows that the strength of the stimuli due to nano-antenna effect has a reduction in comparison with the hot electron injection. We attribute that since the wave vector orthogonality to the plane of the biointerface is important for the nano-antenna effect (Fig. S13 and S14), the refractive index profile change due to the cell membrane and intracellular environment surrounding the metallic nanoislands possibly disturbs the wave vector of the biointerface [28] and decrease the enhancement in the red-spectral window.

## Discussion

Hot-electron injection is the dominant electronic mechanism for the enhancement of capacitive stimulation. The photons both in the blue and red spectral windows have sufficient energy to generate hot electrons. Since the excited state energy of hot electrons under blue excitation is higher than the red excitation, the hot electron injection to the biointerface at blue pump is more probable. But, the decrease in the hot-electron injection probability in the red-spectral region is compensated with the near-field enhancement of the optical energy in the bulk heterojunction (BHJ) composite.

To capture the hot electrons efficiently we formed a Schottky barrier with the PTB7-Th:PC71BM bulk heterojunction (BHJ) composite. Since the generated hot electrons with energies higher than the Schottky barrier energy see lower interfacial resistance toward the BHJ in comparison with the electrode-electrolyte interface, they can be efficiently transferred into the photoactive layer, instead donating the electron to the electrolyte for Faradaic reactions. We also observed this fact from the low magnitude of the Faradaic photocurrents. After the hot electron is injected to the optoelectronic biointerface, the gold nanoislands are left with a hole. Since the capacitive current in the plasmonic biointerface is effectively based on the hole accumulation at the electrode-electrolyte interface, the holes remained after hot electron injection inside the gold nanoislands and the ones that moved to the PTB7-Th due to low potential barrier lead to the capacitive enhancement. Therefore, even though electron relaxation through electron–electron and electron–phonon collisions that are converted into heat energy was generally used as the main mechanism for cell stimulation [4–8], differently in this study both hot electron injection and near-field enhancement are used to improve the displacement currents.

The intensity levels that are applied here for cell stimulation can be safely used for all the tissue or cell types. For example, the most light-sensitive part of human body is the retina, and the design of the biointerfaces need to satisfy maximum permissible exposures for ocular safety limits allowed for ophthalmic applications. In this regard, 1 mW. cm^−2^ light sensitivity of plasmonic biointerface corresponds to more than 3 orders of magnitude smaller than ocular safety limits (see Methods section). Though the transmembrane potential variation in the free membrane seems low, the variation in the attached membrane is significantly larger [27], moreover the current levels (over 100 pA) that are observed by using similar electrophysiological measurements [29] are enough to elicit action potentials. Therefore, these plasmonic biointerfaces are potentially applicable to recover the vision due to photoreceptor loss in the retina [6]. This kind of sight loss is frequent in the clinics due to age-macular degeneration, retinitis pigmentosa, Stargardt's disease [30–32].

Gold nanoislands have high potential for biological applications due to its biocompatibility, easy fabrication and control over optical properties. They have strong absorption from the visible to NIR region, therefore, plasmon-coupled biointerfaces can lead to a new solution for capacitive stimulation for deep-brain regions. The plasmonic absorption arises due to collective oscillation of the conduction-band electrons in the resonance peak of the absorbance spectrum, and its resonance peak of the nanometals can be further controlled by alloying with different metals (e.g., silver) or using different shapes (e.g., cube) [33]. Hence, the combination of the plasmonics with optoelectronic biointerfaces offers a wide variety of device architectures for cell stimulation that can operate at different spectral windows for photomedicine.

In this study, we incorporated plasmonics to light-activated capacitive neurostimulators. Hot-electron injection and nanoantenna effect significantly enhanced the injected charge amounts and currents from the biointerface to the electrolyte. This facilitated variation of membrane potential at single-cell level within the low intensity levels of several mW.cm^−2^, inside the safe limits for retinal photostimulation. In addition to the safe intensity levels, the capacitive charge-injection mechanism is a biocompatible neurostimulation strategy that can be applied long-term and repeatedly to the targeted neurological cell type. We believe that the marriage of plasmonics and optoelectronic biointerfaces point out a bright future for novel nanoengineered optoelectronic neural interfaces.

## Materials and methods

### Materials

Poly-[4,8-bis(5-(2-ethylhexyl)thiophen-2-yl)benzo[1,2-b;4,5-b’]dithiophene-2,6-diyEl-alt-(4-(2-ethylhexyl)-3-fluorothieno[3,4-b]thiophene-)-2-carboxylate-2-6diyl)](PTB7-Th) with a molecular weight of 57,467 g mol^−1^ and [6,6]-PPhenyl-C71-butyric acid methyl ester (PC71BM) with a molecular weight of 1,031 g mol^−1^ were purchased from Ossila Ltd., and used as received. Commercial Gold (Au) with 99.95% purity was used for plasmonic gold nanoisland fabrication. Other materials and processing chemicals such as zinc acetate dihydrate (Zn(CH_3_CO_2_)_2_ ·2H_2_O) with molecular weight 183.48 g mol^−1^, ethanolamine (HOCH_2_CH_2_NH_2_) with molecular weight 61.08 g mol^−1^, and 1,2 dichlorobenzene were purchased from Sigma Aldrich and used without any purification.

### Biointerface preparation

Two type of biointerfaces with and without gold nanoislands (AuNI) were fabricated. Indium Tin Oxide (ITO) on glass substrates (side length, 20 mm; side length, 15 mm; thickness 1.1 mm; resistance, 14 - 16 Ω.cm^−2^; Ossila) was cleaned with 10 wt% Sodium Hydroxide solution for 5 min, 2 % by volume a specific tension-active agent in a water solution (HELLMANEX II, 3 %) for 15 mins at 55 °C, DI water for 15 min, acetone for 15 mins and isopropyl alcohol for 15 mins using ultrasonic bath. The cleaned ITO on glass coated substrates were dried in oven at 50°C and treated with UV-Ozone for 15 min. ZnO precursor sol-gel solution (0.45 M) was prepared by mixing 219.3 mg zinc acetate dehydrate (Zn(CH₃CO₂)₂·2H₂O), 2 ml of 2-Methoxyethanol (C_3_H_8_O_2_) and 73 mg of Ethanolamine (HOCH₂CH₂NH₂) and ultrasonicating at 55°C for 15 min [24]. 14.4 mg ml^−1^ of the donor co-polymer PTB7-Th and 21.6 mg ml^−1^ of acceptor molecule PC71BM solution was prepared in o-dichlorobenzene and stirred overnight at 70°C on a hot plate (niye o sayilar). Then 14.4 mg ml^−1^ PTB7-Th, 21.6 mg ml^−1^ PC71BM (1:1.5 wt. ratio) in o-dichlorobenzene and 3 % (in volume) 1,8-Diiodooctane (C8H16I2) was added and stirred for 3 h on a hot plate at 70°C. The control biointerface was fabricated by spin coating ZnO precursor sol-gel solution at 2000 rpm for 60 s on a cleaned ITO coated on glass substrate and annealed on a hot plate at 290°C for 15 min [22]. Further on ITO/ZnO substrate photoactive blend of PTB7-Th:PC71BM (1:1.5 wt. ratio) solution in o-dichlorobenzene was spin coated at 600 rpm for 200 s, then dried under nitrogen flow for 15 min and heated at 100 °C for 5 min. Plasmonic biointerface was fabricated on a control biointerface by coating 5 nm of Au using thermal evaporator (Bruker, detail) at the rate of 0.02-0.03 nm / s under 6.0 × 10^−6^ mbar vacuum pressure. Due to thin layer of gold forming nanoislands even at room temperature, it is annealed at 100°C, which is maximum temperature that photoactive layer can withstand, before its optical properties deteriorating.

### Optical and surface characterization

UV-Vis Transmittance (Edinburg) was used to measure absorbance of the control and plasmonic biointerfaces. Scanning electron microscopy (SEM) (Zeiss) was used for imaging surface profile and the cross-sectional structure of the control and plasmonic biointerfaces. For better visualization of the surface an additional thin gold coating is deposited.

### Photocurrent measurements

Further, the photocurrent measurements were performed on a set up including an Olympus T2 upright microscope. An extracellular patch clamp (EPC) 800 patch clamp amplifier (HEKA Elektronik) is used for measuring photocapacitive currents. The experiment was carried out at room temperature in an extracellular aCSF aqueous medium as supporting electrolyte solution. Extracellular medium (aCSF) was prepared by mixing 10 mM of 4-(2-hydroxyethyl)-1-piperazineethanesulfonic acid (HEPES), 10 mM of glucose, 2 mM CaCl2, 140 mM of NaCl, 1 mM of MgCl2, 3 mM of KCl, into the distilled water and pH was calibrated to 7.4 using 1M NaOH. For the light stimulation, Thorlab’s blue LED (M450LP1) with nominal wavelength of 445 nm was varied between 165 μW. cm^−2^ – 169 mW. cm^−2^, green LED (M530L3) with nominal wavelength of 530 nm was varied between 89 μW. cm^−2^ – 41 mW. cm^−2^ and with nominal wavelength of 630 nm was varied between 64 μW. cm^−2^ – 85 mW. cm^−2^ (Fig. S2, S3). All LEDs were driven by DC2200 - High-Power 1-Channel LED Driver with Pulse Modulation (Thorlab’s). A power meter (Newport 843-R) was used to measure the exact power of light reaching the surface of biointerface. The illumination was focused at water immersion objective holder from the photoactive layer side of the biointerface. Illumination area was 1 cm^2^. The biointerface corners were cleaned with toluene to stabilize the photocurrent measurements.

### Electrochemical measurements

The electrochemical impedance spectroscopy (EIS) in frequency response analysis (FRA) potential scan mode were performed on an Autolab Potentiostat Galvanostat PGSTAT (Metrhom, Netherlands). A three-electrode setup consisting of Ag/AgCl as the reference electrode, platinum wire as the counter electrode and the biointerfaces as working electrode were used. The experiment was carried out at room temperature in an extracellular aCSF aqueous medium as supporting electrolyte solution. The biointerfaces were excited by blue, green and red LEDs with optical power output of 25 mW.cm^−2^. In this mode, a sinusoidal alternating voltage perturbation with root mean square value of 10 mV was applied while the frequency was varied from 1 Hz to 0.1 MHz. The electrochemical responses were fitted with proper circuit diagram using NOVA software.

Using the macro-electrical parameters obtained by fitting, the driving electrical parameters in the bulk of the biointerface such as effective lifetime (*τ*_*n*_) and mobility (*μ_n_*) of charge carriers can be estimated using following equations;

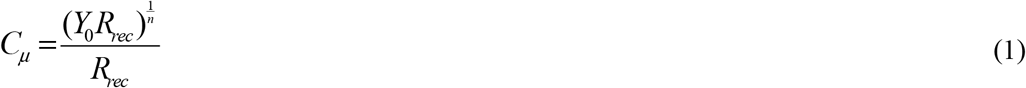

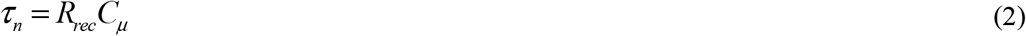

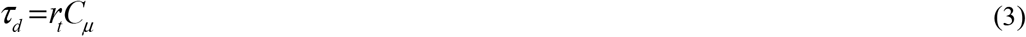

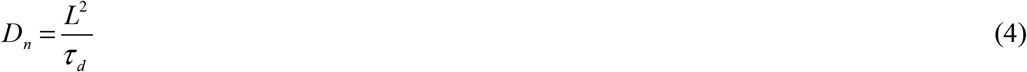

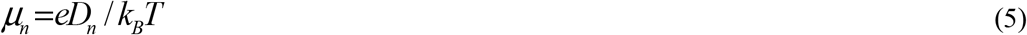

Where Cμ the chemical capacitance corresponds to carrier accumulation in the bulk, *τ*_*d*_ the electron diffusion time, *D*_*n*_ the diffusion coefficient of electrons, *L* the photoactive thin film thickness, and *k*_*B*_*T* the thermal energy.

### MTT Assay

Cytotoxic effect of biointerfaces on SHSY-5Y cells were assessed by measuring the mitochondrial activity with MTT assay. MTT assay was performed as described by Bahmani et al. [7]. Briefly, biointerfaces and ITO control were sterilized with 70% ethanol and followed by UV-C treatment for 5 min. Sterilized samples were placed in a 6-well cell culture plate. Then, SH-SY5Y with 0.3 × 10^6^ cells were seeded to each well in Dulbecco's Modified Eagle Medium (DMEM) containing 10% fetal bovine serum (FBS) and 1% penicillin-streptomycin. Cell attachment and growth were allowed by 48h incubation at 37°C and 5% CO_2_. After incubation, growth medium was discarded, and samples were transferred to a new 6-well plate. 1 mg/ml 3-(4,5-Dimethylthiazol-2-yl)-2,5-diphenyltetrazolium bromide (MTT) was added onto the samples in serum-free DMEM and incubated for 4h at 37°C to allow formazan formation. MTT solution was removed and formazan salts were dissolved in 50:50 (v:v) ethanol/dimethyl sulfoxide mixture (EtOH/DMSO). Absorbance of each sample and ITO control were measured in a 96-well plate with Synergy H1 microplate reader (BioTek). Relative absorbance of biointerface samples to ITO was calculated to determine relative biocompatibility.

### LDH Leakage Assay

LDH leakage assay (CytoSelect LDH cytotoxicity assay kit, CBA-241, Cell Biolabs) was performed to assess the membrane integrity of the cells grown on the substrates. Experiment was performed according to manufacturer’s manual. Briefly, 3 × 10^5^ SHSY-5Y cells were seeded onto sterilized biointerfaces and control groups as explained in cytotoxicity assessment section. Two sets of ITO control were included as positive and negative control groups. Samples were incubated at 37°C and 5% CO_2_ in a time dependent manner (24h, 48h and 72h). Negative and positive ITO controls were treated with 1% Triton X-100 and dH2O, respectively. 90 μL medium from each sample was transferred to a 96-well plate and mixed with 10 μL LDH cytotoxicity assay reagent. The plate was incubated at 37°C for 30min. Presence of LDH in the medium was measured by absorbance at 450 nm with Synergy H1 microplate reader (BioTek).

### Immunolabeling and Fluorescence Microscopy

Substrates were sterilized and SHSY-5Y cells were seeded on substrates in a 6-well plate as explained in cytotoxicity assessment section. The 6-well plate was incubated at 37°C and 5% CO2 for 48h. Cells grown on the substrates were washed three times with phosphate-buffered saline containing 0.1% Triton X-100 (PBSt). Cell fixation was performed by 30min incubation at 37°C with 4% paraformaldehyde (PFA). Fixed cells were washed three times with PBSt, and blocked with PBSt containing 5% bovine serum albumin (BSA) for 2h at RT. Anti-beta III tubulin (ab78078, Abcam) primary antibody was used for cytoskeleton labelling and 4’,6-Diamidino-2-Phenylindole (DAPI) staining was performed for nucleus visualization. Samples were incubated with 1:250 diluted Anti-beta III tubulin primary antibody for 2h, and washed three times with PBSt. Alexa Fluor conjugated goat anti-mouse IgG H&L (ab150113) secondary antibody (1:1000) and DAPI (1:10000) was diluted in PBSt containg 5% BSA and samples were incubated in the mixture for 1h. Substrates washed three times with PBSt and mounted onto microscope slides with Mowiol. Imaging was performed with a fluorescence microscope (Zeiss-ObserverZ1).

### Electrophysiology

Electrophysiology measurements were performed with EPC 800 patch clamp amplifier (HEKA Elektronik). The biointerfaces were cleaned with 70% (by volume) ethanol solution and incubated for 2 day in water. The pulled patch pipettes of 8-12 MΩ were used to perform the whole-cell patch clamp experiment. Extracellular medium (aCSF) was prepared as previous mentioned. Internal cellular medium was prepared by mixing 140 mM KCl, 2 mM MgCl_2_, 10 mM HEPE, 10 mM ethylene glycol-bis(β-aminoethyl ether)−N,N,N’,N’-tetraacetic acid (EGTA), 2 mM Mg-ATP in water and its pH was calibrated to 7.2-7.3 using 1 M KOH, and patch pipettes were filled with intracellular solution. Digital camera integrated with Olympus T2 upright microscope was used to monitor the cells. The whole-cell patched cells were investigated up to 30 mins to avoid damage done by patched pipettes.

### Optical safety considerations

According to ocular safety standards for ophthalmic applications, the maximum permissible radiant power (MPF) that could enter the pupil chronically or in a single short exposure is calculated. For single-pulse exposures (pulse width of 10ms) at 400 - 700 nm, the peak limit is governed by the equation MPF = 6.93 × 10^−4^ C_T_C_E_t^−0.25^, where, t = 10 ms (pulse width), C_T_ × 1 in the range of 400–700 nm and C_E_ is a function of the visual angle α. For retinal spot sizes greater than 1.7 mm, α = α_max_ = 100 mrad and C_E_ = 6.67 × 10^−3^ α^2^ = 66.7 W. For 10 ms pulse width MPF = 146.17 mW. For a pupil size with diameter 1.7 mm MPF per unit area is MPF_per unit area_ = 6443 mW. cm^−2^.

## Acknowledgement

This project has received funding from the European Research Council (ERC) under the European Union Horizon 2020 Research and Innovation Programme (Grant agreement no. 639846). SN acknowledges the support by the Turkish Academy of Sciences (TÜBA-GEBİP; The Young Scientist Award Program) and the Science Academy of Turkey (BAGEP; The Young Scientist Award Program). The authors thank to Dr. Arif Engin Cetin in Izmir Biomedicine and Genome Center, and Dr. Gokhan Demirel in Gazi University for the fruitful discussions.

## Contributions

R.M. S.S and S.N. designed the experiments. U.M.D and I.H.K performed cell seeding and biological assays. R.M. S.S, and O.K. performed the electrophysiology and photocurrent experiments. R.M. and S.S performed the electrochemical measurements. I.B.D designed optical setup. S.S and H.B.J helped R.M. for optical measurements. R.M, S.S, B.U and S.N. analysed the data. R.M., S.S, O.K and S.N. wrote the manuscript, with input from all authors.

## References

1. da Cruz, L., et al., Five-year safety and performance results from the Argus II retinal prosthesis system clinical trial. Ophthalmology, 2016. 123(10): p. 2248–2254.

2. Yuan, X., A. Hierlemann, and U. Frey, Dual-mode microelectrode array with 20k-electrodes and high SNR for high-throughput extracellular recording and stimulation. Frontiers in Cellular Neuroscience, 2018. 12.

3. Grill, W.M. and C.C. Mclntyre, Extracellular excitation of central neurons: implications for the mechanisms of deep brain stimulation. Thalamus & Related Systems, 2001. 1(3): p. 269–277.

4. Bareket-Keren, L. and Y. Hanein, Novel interfaces for light directed neuronal stimulation: advances and challenges. International journal of nanomedicine, 2014. 9(Suppl 1): p. 65.

5. Rand, D., et al., Direct Electrical Neurostimulation with Organic Pigment Photocapacitors. Advanced Materials, 2018. 30(25): p. 1707292.

6. Ghezzi, D., et al., A polymer optoelectronic interface restores light sensitivity in blind rat retinas. Nature Photonics, 2013. 7(5): p. 400.

7. Bahmani Jalali, H., et al., Effective neural photostimulation using indium-Based type-II quantum dots. ACS nano, 2018. 12(8): p. 8104–8114.

8. Jiang, Y., et al., Rational design of silicon structures for optically controlled multiscale biointerfaces. Nature biomedical engineering, 2018. 2(7): p. 508.

9. Gilbert, F., A.R. Harris, and R.M. Kapsa, Controlling brain cells with light: ethical considerations for optogenetic clinical trials. AJOB Neuroscience, 2014. 5(3): p. 3–11.

10. Maya-Vetencourt, J.F., et al., A fully organic retinal prosthesis restores vision in a rat model of degenerative blindness. Nature materials, 2017. 16(6): p. 681.

11. Di Maria, F., et al., The evolution of artificial light actuators in living systems: from planar to nanostructured interfaces. Chemical Society Reviews, 2018. 47(13): p. 4757–4780.

12. Zimmerman, J.F. and B. Tian, Nongenetic optical methods for measuring and modulating neuronal response. ACS nano, 2018. 12(5): p. 4086–4095.

13. Tatsuma, T., H. Nishi, and T. Ishida, Plasmon-induced charge separation: chemistry and wide applications. Chemical science, 2017. 8(5): p. 3325–3337.

14. Carvalho-de-Souza, J.L., et al., Photosensitivity of neurons enabled by cell-targeted gold nanoparticles. Neuron, 2015. 86(1): p. 207–217.

15. Eom, K., et al., Enhanced infrared neural stimulation using localized surface plasmon resonance of gold nanorods. Small, 2014. 10(19): p. 3853–3857.

16. Yoo, S., et al., Photothermal inhibition of neural activity with near-infrared-sensitive nanotransducers. ACS nano, 2014. 8(8): p. 8040–8049.

17. Lee, J.W., et al., Gold nanostar-mediated neural activity control using plasmonic photothermal effects. Biomaterials, 2018. 153: p. 59–69.

18. Nakatsuji, H., et al., Thermosensitive Ion Channel Activation in Single Neuronal Cells by Using Surface-Engineered Plasmonic Nanoparticles. Angewandte Chemie International Edition, 2015. 54(40): p. 11725–11729.

19. Soganci, I.M., et al., Localized plasmon-engineered spontaneous emission of CdSe/ZnS nanocrystals closely-packed in the proximity of Ag nanoisland films for controlling emission linewidth, peak, and intensity. Optics express, 2007. 15(22): p. 14289–14298.

20. Hoang, C.V., et al., Interplay of hot electrons from localized and propagating plasmons. Nature communications, 2017. 8(1): p. 771.

21. Zhao, L., et al., Two effects of 1, 8-diiodooctane on PTB7-Th: PC71BM polymer solar cells. Organic Electronics, 2016. 34: p. 188–192.

22. Srivastava, S.B., et al., Band Alignment Engineers Faradaic and Capacitive Photostimulation of Neurons Without Surface Modification. Physical Review Applied, 2019. 11(4): p. 044012.

23. Bhattacharya, G., et al., Probing the flat band potential and effective electronic carrier density in vertically aligned nitrogen doped diamond nanorods via electrochemical method. Electrochimica Acta, 2017. 246: p. 68–74.

24. Srivastava, S.B., P. Sonar, and S.P. Singh, Charge transport studies in donor-acceptor block copolymer PDPP-TNT and PC71BM based inverted organic photovoltaic devices processed in room conditions. AIP Advances, 2015. 5(7): p. 077177.

25. Maurano, A., et al., Recombination dynamics as a key determinant of open circuit voltage in organic bulk heterojunction solar cells: a comparison of four different donor polymers. Advanced Materials, 2010. 22(44): p. 4987–4992.

26. Abdullaeva, O.S., et al., Organic Photovoltaic Sensors for Photocapacitive Stimulation of Voltage-Gated Ion Channels in Neuroblastoma Cells. Advanced Functional Materials, 2019. 29(21): p. 1805177.

27. Jakešová, M., et al., Optoelectronic control of single cells using organic photocapacitors. Science advances, 2019. 5(4): p. eaav5265.

28. Gather, M.C. and S.H. Yun, Single-cell biological lasers. Nature Photonics, 2011. 5(7): p. 406.

29. Parameswaran, R., et al., Photoelectrochemical modulation of neuronal activity with free-standing coaxial silicon nanowires. Nature nanotechnology, 2018. 13(3): p. 260.

30. Bressler, N.M., S.B. Bressler, and S.L. Fine, Age-related macular degeneration. Survey of ophthalmology, 1988. 32(6): p. 375–413.

31. Hartong, D.T., E.L. Berson, and T.P. Dryja, Retinitis pigmentosa. The Lancet, 2006. 368(9549): p. 1795–1809.

32. Delori, F.C., et al., In vivo measurement of lipofuscin in Stargardt’s disease--Fundus flavimaculatus. Investigative ophthalmology & visual science, 1995. 36(11): p. 2327–2331.

33. Sau, T.K. and A.L. Rogach, Complex-shaped metal nanoparticles. 2012: Wiley Online Library.

