## supplementary figures for "Plasmon-Coupled Photocapacitor Neuromodulators"

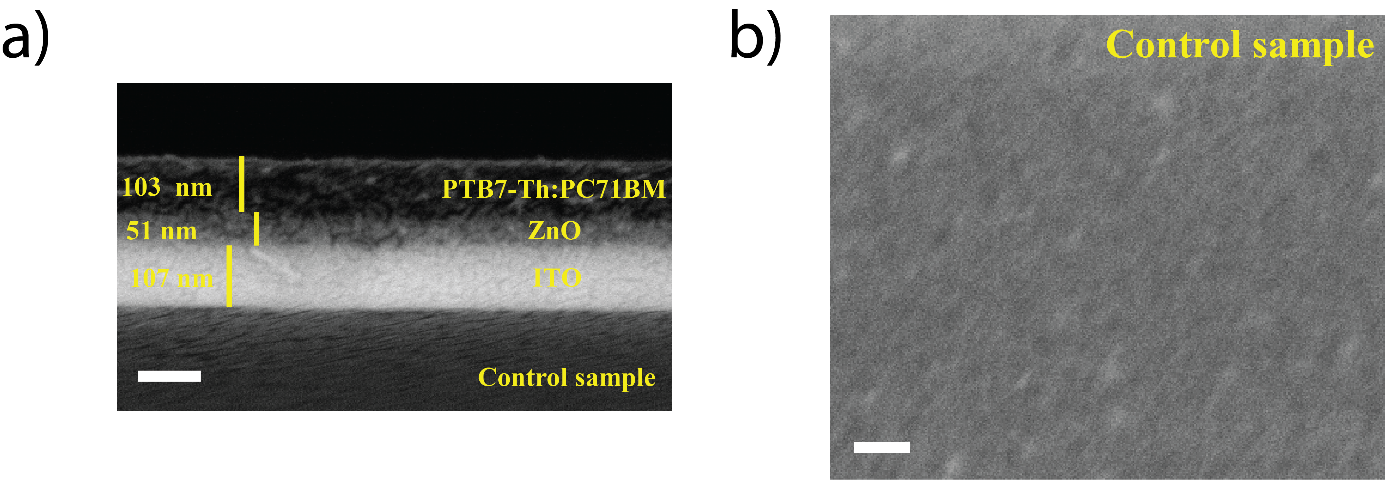

Figure S1. (a) Cross-sectional scanning electron microscopy image of the control biointerface (scale bar, 100 nm). (b) Scanning electron microscopy image of the control biointerface surface (scale bar, 100 nm).

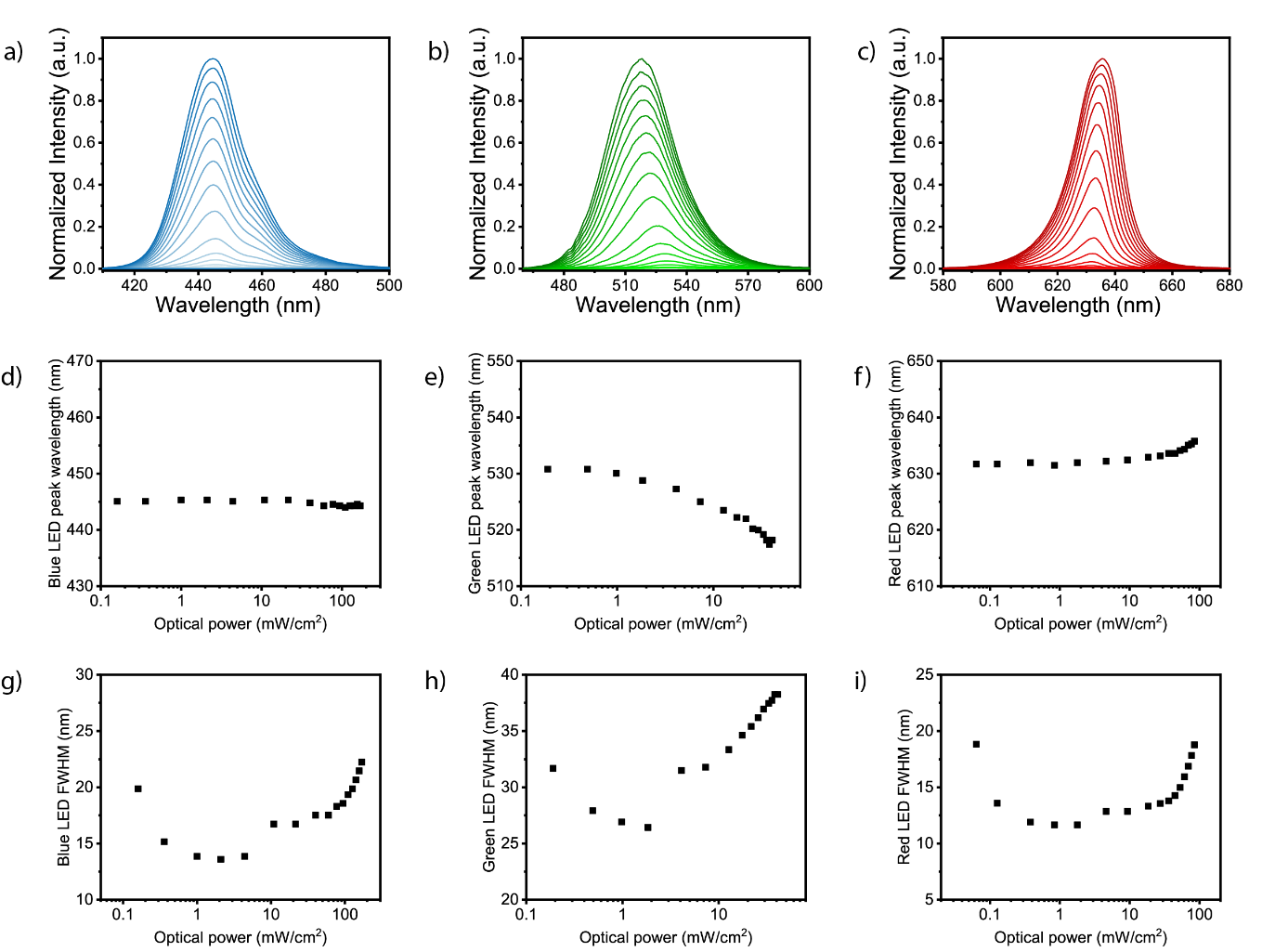

Figure S2. (a) – (c) Spectral intensity, (d) – (f) peak wavelength and (g) – (i) FWHM properties of blue (a,d,g), green (b,e,h) and red (c,f,i) illumination sources at different optical powers.

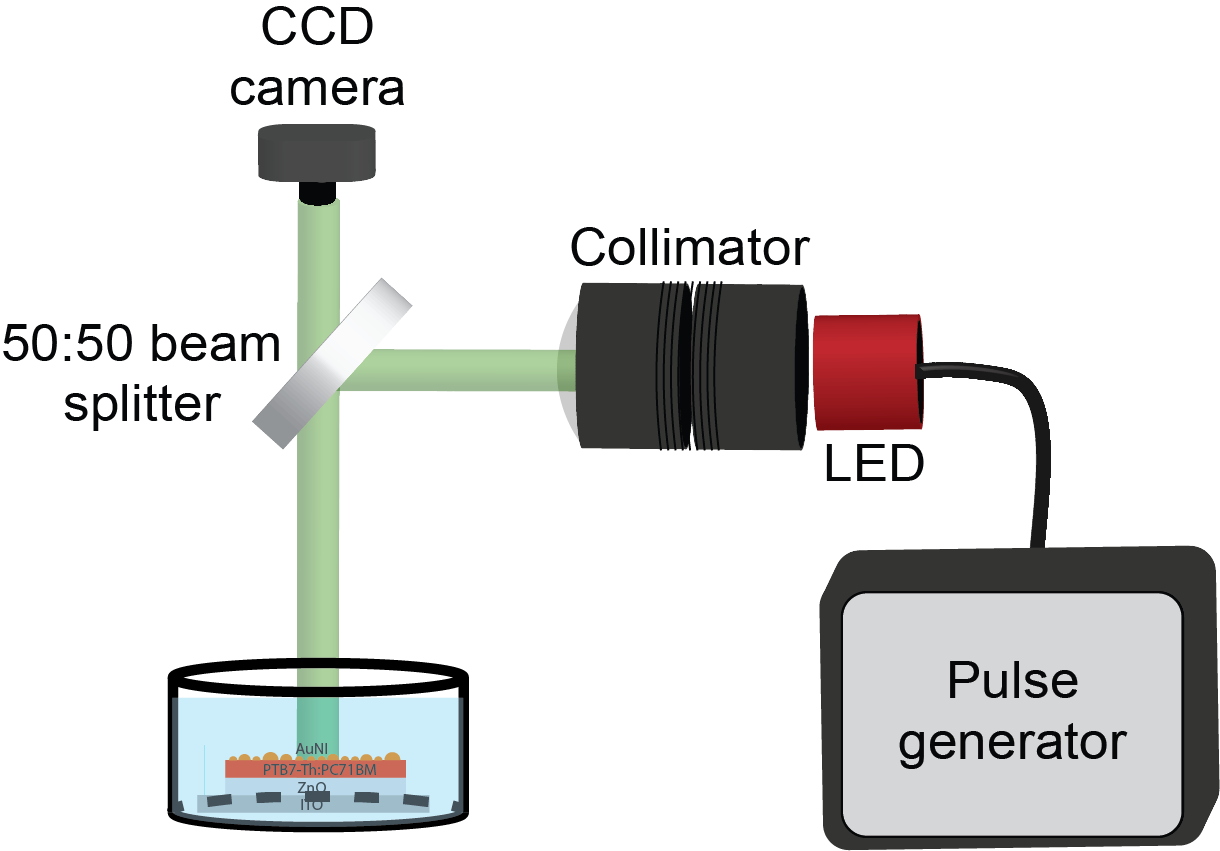

Figure S3. Optical set-up during photocurrent and electrophysiology experiments.

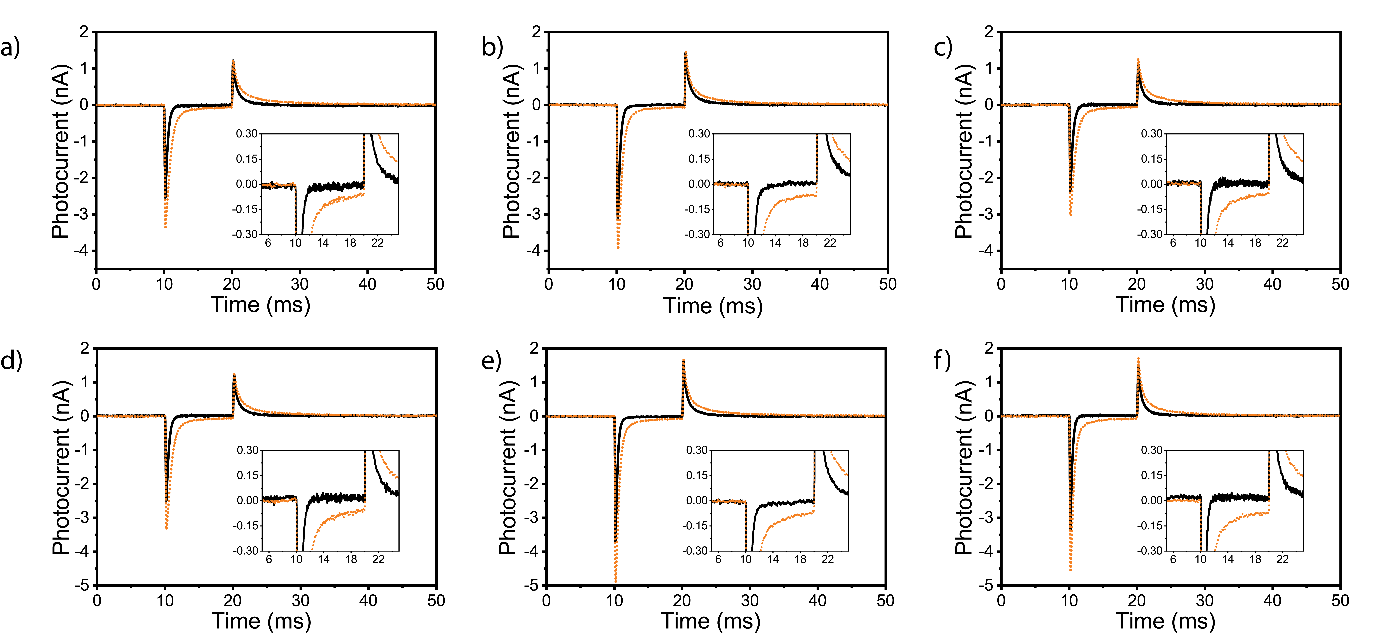

Figure S4. Photocurrents generated by the control and plasmonic biointerfaces under free standing mode. (a) – (c) Photocurrent vs time for control (black line) and plasmonic (orange dotted line) biointerfaces illuminated under same photon counts. The inset zooms low photocurrent levels. All axes are kept equal. (d) – (f) Photocurrent vs time for control (black line) and plasmonic (orange dotted line) biointerfaces excited under same optical power. The inset zooms low photocurrent levels. All axes are kept equal. ((a,d) – blue light illumination, (b,e) – green light illumination and (c,f) – red light illumination).

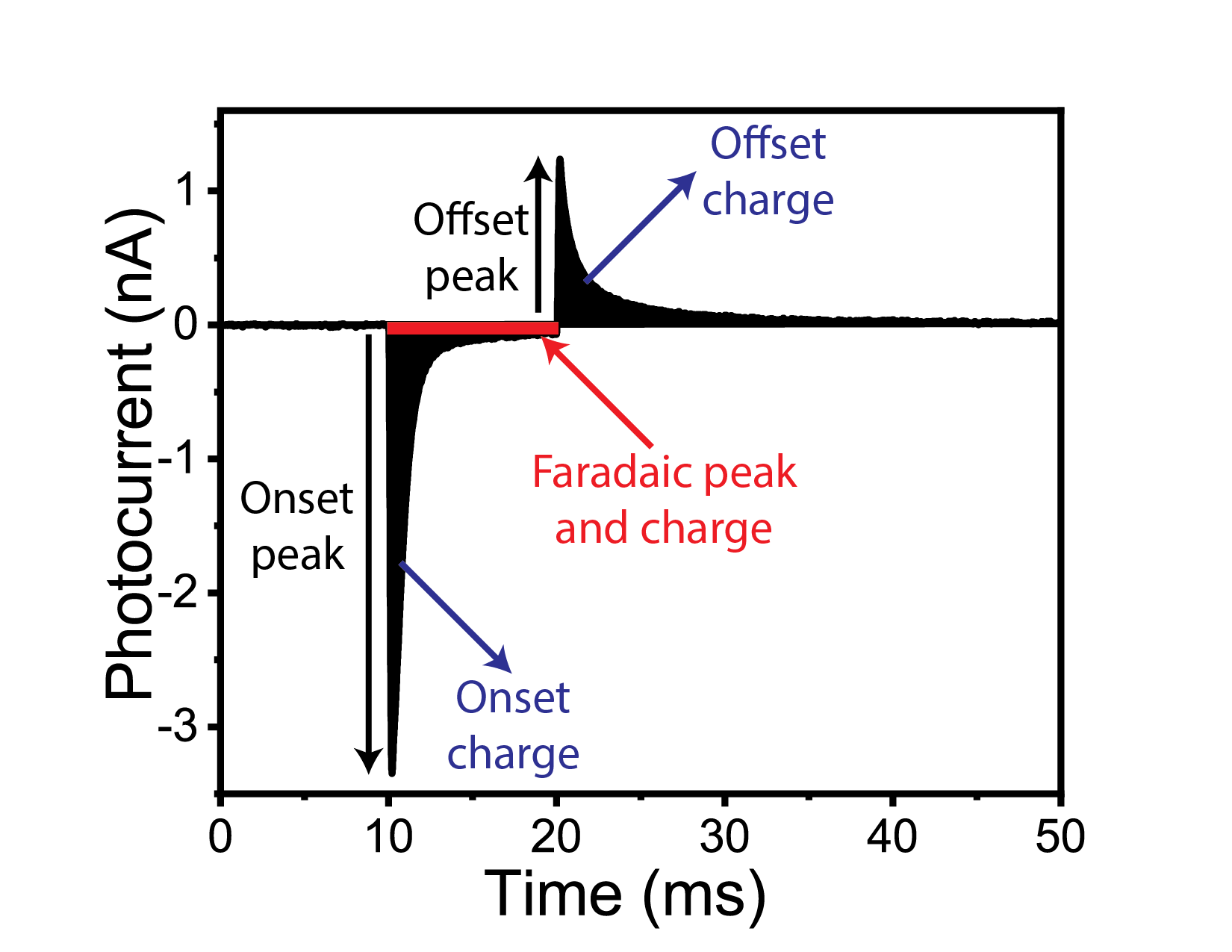

Figure S5. Peaks and charges of capacitive and faradaic photocurrent components.

$Enhancement in onset peak=100\times(\frac{plasmonic onset peak-control onset peak}{control onset peak} )$	(S1)

$Enhancement in onset charge=100\times(\frac{plasmonic onset charge-control onsetcharge}{control onset charge} )$	(S2)

$Faradaic peak percentage=100\times(\frac{Faradaic peak}{onset peak} )$	(S3)

$Faradaic charge percentage=100\times(\frac{Faradaic charge}{onset charge} )$	(S4)

Table S1. Onset charge amount, Faradaic charge amount, Faradaic charge percentage, enhancement in charge for both control and plasmonic biointerfaces under blue, green and red light illumination having same photon counts (8.8±0.2 x 10^16^ photons s^-1^.cm^-2^).

|  | Blue | | Green | | Red | |
| --- | --- | --- | --- | --- | --- | --- |
|  | Control | Plasmonic | Control | Plasmonic | Control | Plasmonic |
| Onset charge (pC) | 6.95 | 19.85 | 7.75 | 20.40 | 5.70 | 17.55 |
| Faradaic charge (pC) | 0.02 | 0.05 | 0 | 0.05 | 0 | 0.04 |
| Faradaic charge (%) | 0.1 | 0.25 | 0 | 0.24 | 0 | 0.24 |
| Enhancement in onset charge (%) | 185.6 | | 163.2 | | 207.9 | |

Table S2. Onset charge amount, Faradaic charge amount, Faradaic charge percentage, enhancement in charge for both control and plasmonic biointerfaces under blue, green and red light illumination having same optical power (42 ± 2 mW.cm^-2^).

|  | Optical power | | | | | |
| --- | --- | --- | --- | --- | --- | --- |
|  | Blue | | Green | | Red | |
|  | Control | Plasmonic | Control | Plasmonic | Control | Plasmonic |
| Onset charge (pC) | 6.95 | 19.85 | 8.38 | 22.45 | 6.77 | 20.45 |
| Faradaic charge (pC) | 0.02 | 0.05 | 0 | 0.06 | 0 | 0.06 |
| Faradaic charge (%) | 0.1 | 0.25 | 0 | 0.26 | 0 | 0.29 |
| Enhancement in onset charge (%) | 185.6 | | 167.9 | | 202.0 | |

Table S3. Onset magnitude, Faradaic magnitude, Faradaic magnitude percentage, enhancement in magnitude for both control and plasmonic biointerfaces under blue, green and red light illumination having same photon counts (8.8±0.2 x 10^16^ photons s^-1^.cm^-2^).

|  | Counts | | | | | |
| --- | --- | --- | --- | --- | --- | --- |
|  | Blue | | Green | | Red | |
|  | Control | Plasmonic | Control | Plasmonic | Control | Plasmonic |
| Onset peak (nA) | 2.49 | 3.35 | 3.15 | 3.96 | 2.36 | 3.02 |
| Faradaic peak (nA) | 0.010 | 0.022 | 0 | 0.021 | 0 | 0.006 |
| Faradaic peak (%) | 0.40 | 0.65 | 0 | 0.53 | 0 | 0.35 |
| Enhancement in onset peak (%) | 34.5 | | 25.7 | | 28.0 | |

Table S4. Onset magnitude, Faradaic magnitude, Faradaic magnitude percentage, enhancement in magnitude for both control and plasmonic biointerfaces under blue, green and red light illumination having same optical power (42 ± 2 mW.cm^-2^).

|  | Optical power | | | | | |
| --- | --- | --- | --- | --- | --- | --- |
|  | Blue | | Green | | Red | |
|  | Control | Plasmonic | Control | Plasmonic | Control | Plasmonic |
| Onset peak (nA) | 2.49 | 3.35 | 3.70 | 4.89 | 3.37 | 4.55 |
| Faradaic peak (nA) | 0.010 | 0.022 | 0 | 0.026 | 0 | 0.016 |
| Faradaic peak (%) | 0.40 | 0.65 | 0 | 0.06 | 0 | 0.35 |
| Enhancement in onset peak (%) | 34.5 | | 32.2 | | 35.0 | |

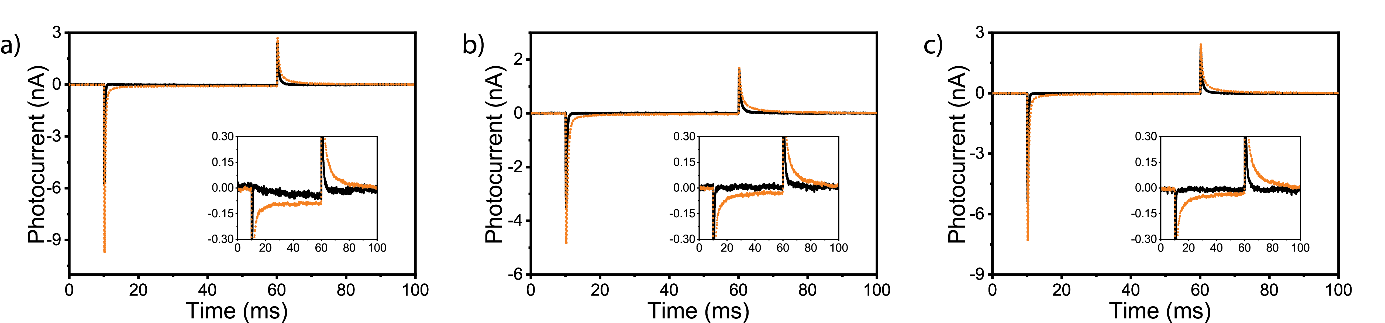

Figure S6. (a) – (c) Photocurrent vs time for control (black line) and plasmonic (orange line) biointerfaces illuminated under blue (a), green (b) and red (c) light with 50 ms pulse width and 100 ms period. The inset zooms low photocurrent level.

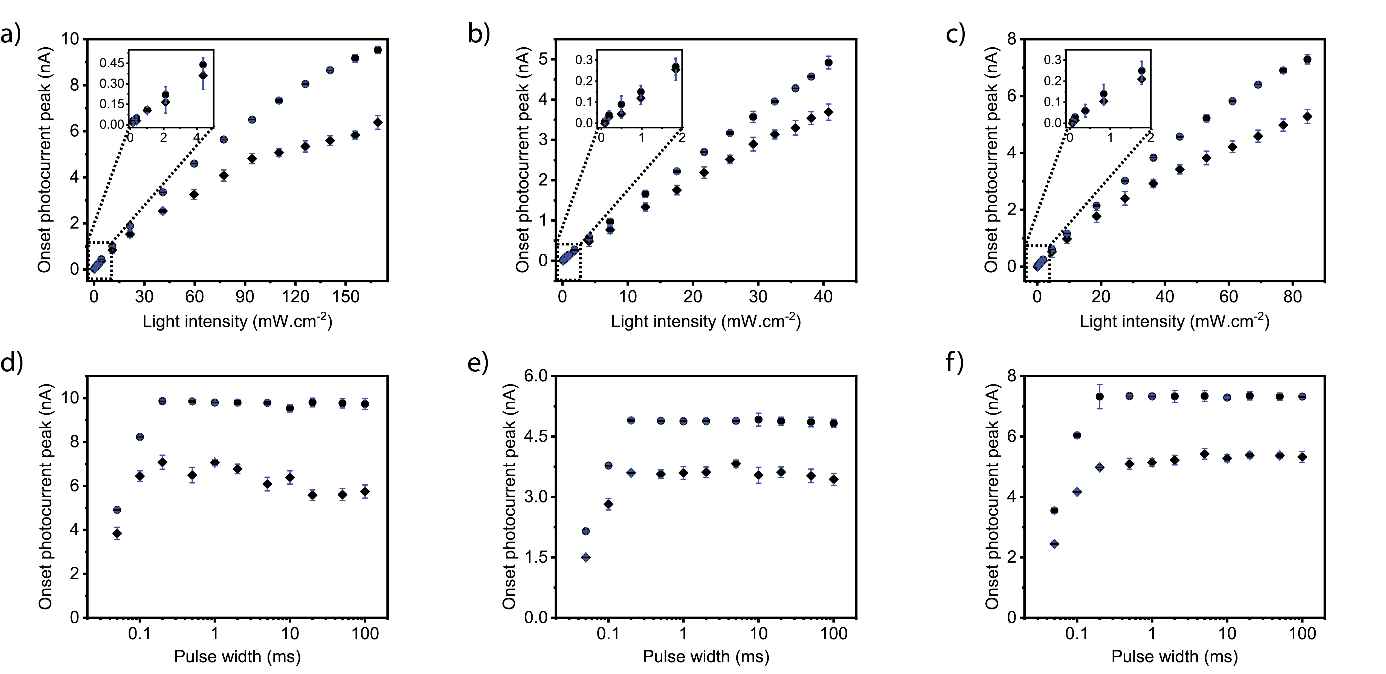

Figure S7. (a) - (c) Magnitude of onset peak of capacitive photocurrents of control (rhombus dots) and plasmonic (circle dots) biointerface as a function of light intensity. The inset zooms low light intensity levels. (d) - (f) Magnitude of onset peak of capacitive photocurrent of control (rhombus dots) and plasmonic (circle dots) biointerface as a function of pulse width under maximum light intensity. ((a,d) – blue light illumination, (b,e) – green light illumination and (c,f) – red light illumination.)

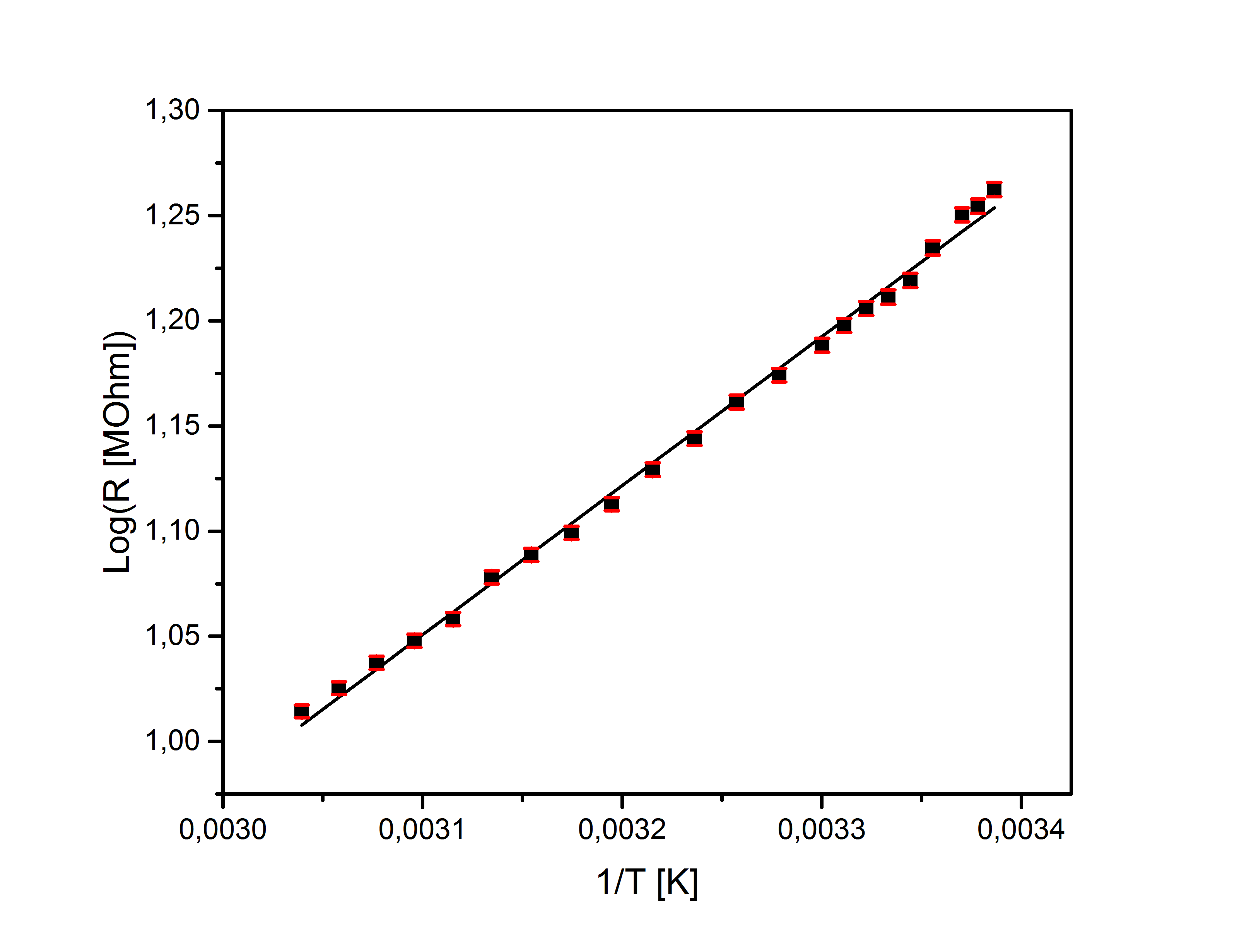

Figure S8. Temperature dependence of the pipette tip resistance. The pipette resistance did not change under continuous light illumination at 170 mW.cm^-2^.

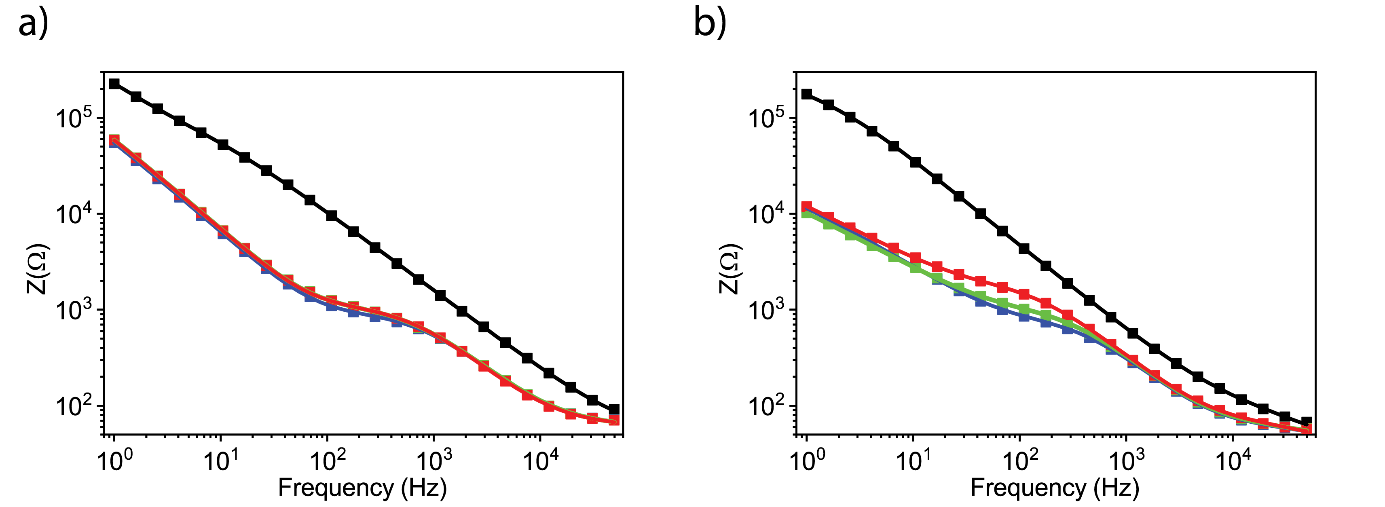

Figure S9. Electrochemical impedance spectroscopy of control and plasmonic biointerfaces under dark, blue, green and red light illumination at zero dc bias. (a) Bode magnitude of control biointerface. (b) Bode magnitude of plasmonic biointerface. (Squares - experimental data, lines – fitted data, black – no illumination, blue – blue light illumination, green – green light illumination and red – red light illumination)

Table S5. Fitted electrical parameters of the impedance analysis.

| SAMPLE | Light | R_s_(Ω) | R_rec_(kΩ) | CPE1 | W | C_dl_(µF) | R_l_(Ω) | C_g_(nF) | χ^2^(%) |
| --- | --- | --- | --- | --- | --- | --- | --- | --- | --- |
| Plasmonic | Dark | 45 | 26.300 | 3.50, 0.716 | 3.14 | 0.44 | 91.4 | 82.3 | 1.13 |
| Plasmonic | Blue | 45 | 0.663 | 1.14, 0.886 | 30.10 | 3.14 | 31.3 | 146.0 | 1.07 |
| Plasmonic | Green | 45 | 0.868 | 1.14, 0.882 | 35.80 | 3.06 | 33.7 | 137.0 | 1.10 |
| Plasmonic | Red | 45 | 1.610 | 1.32, 0.861 | 30.60 | 3.10 | 43.1 | 148.0 | 1.17 |
| Control | Dark | 45 | 72.600 | 0.84, 0.726 | 0.89 | 0.51 | 76.5 | 35.6 | 0.81 |
| Control | Blue | 45 | 0.777 | 0.50, 0.906 | 1.95 | 2.28 | 30.7 | 41.6 | 1.39 |
| Control | Green | 45 | 0.923 | 0.67, 0.880 | 1.82 | 2.13 | 24.7 | 31.3 | 3.10 |
| Control | Red | 45 | 0.919 | 0.57, 0.894 | 1.80 | 2.14 | 29.7 | 42.3 | 1.40 |

Table S6. Mobility and effective lifetime estimation using impedance analysis.

| SAMPLE | Light | C_µ_(µF) | τ_d_(µs) | τ_n_(µs) | D_n_(nm) | µ_n_(cm^2^/Vs)(x10^-5^) |
| --- | --- | --- | --- | --- | --- | --- |
| Plasmonic | D | 1.36 | 61.14 | 35700.00 | 0.16 | 0.63 |
| Plasmonic | B | 0.45 | 20.16 | 297.02 | 0.50 | 1.92 |
| Plasmonic | G | 0.45 | 20.33 | 392.15 | 0.49 | 1.90 |
| Plasmonic | R | 0.49 | 21.99 | 786.93 | 0.46 | 1.76 |
| Control | D | 0.29 | 13.22 | 21300.00 | 0.80 | 3.10 |
| Control | B | 0.22 | 9.96 | 172.00 | 1.06 | 3.73 |
| Control | G | 0.24 | 10.99 | 225.40 | 0.96 | 3.91 |
| Control | R | 0.23 | 10.47 | 213.89 | 1.01 | 3.10 |

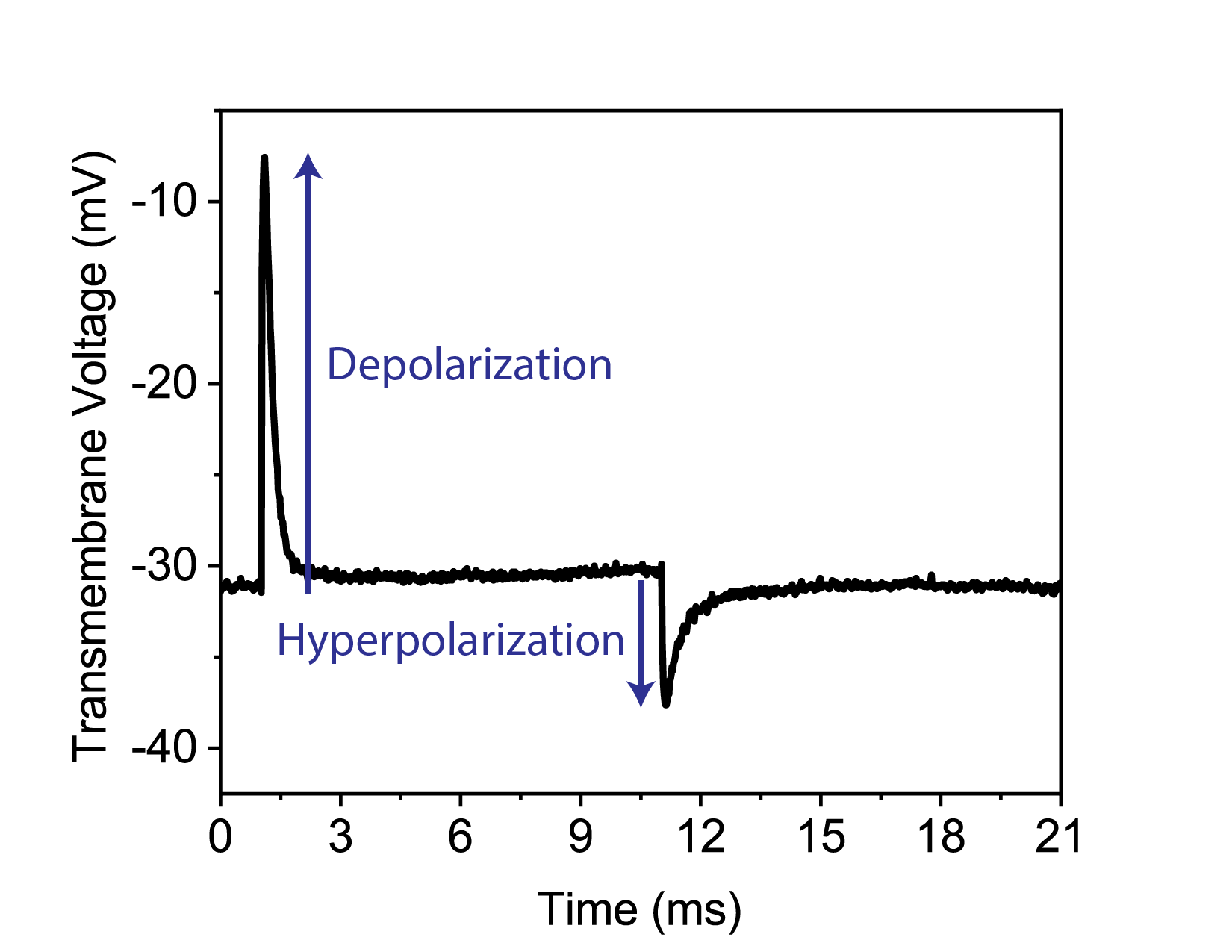

Figure S10. Transmembrane potential variation of SHSY-5Y cell on the plasmonic biointerface under blue LED excitation with 10 ms pulse width.

$Enhancement in depolarization=100\times(\frac{plasmonic depolarization-control depolarization}{control depolarization} )$	(s5)

$Overall enhancement in depolarization=\sum_{optical power/frequency}^{N} \frac{Enhancement in depolarization}{N}$	(s6)

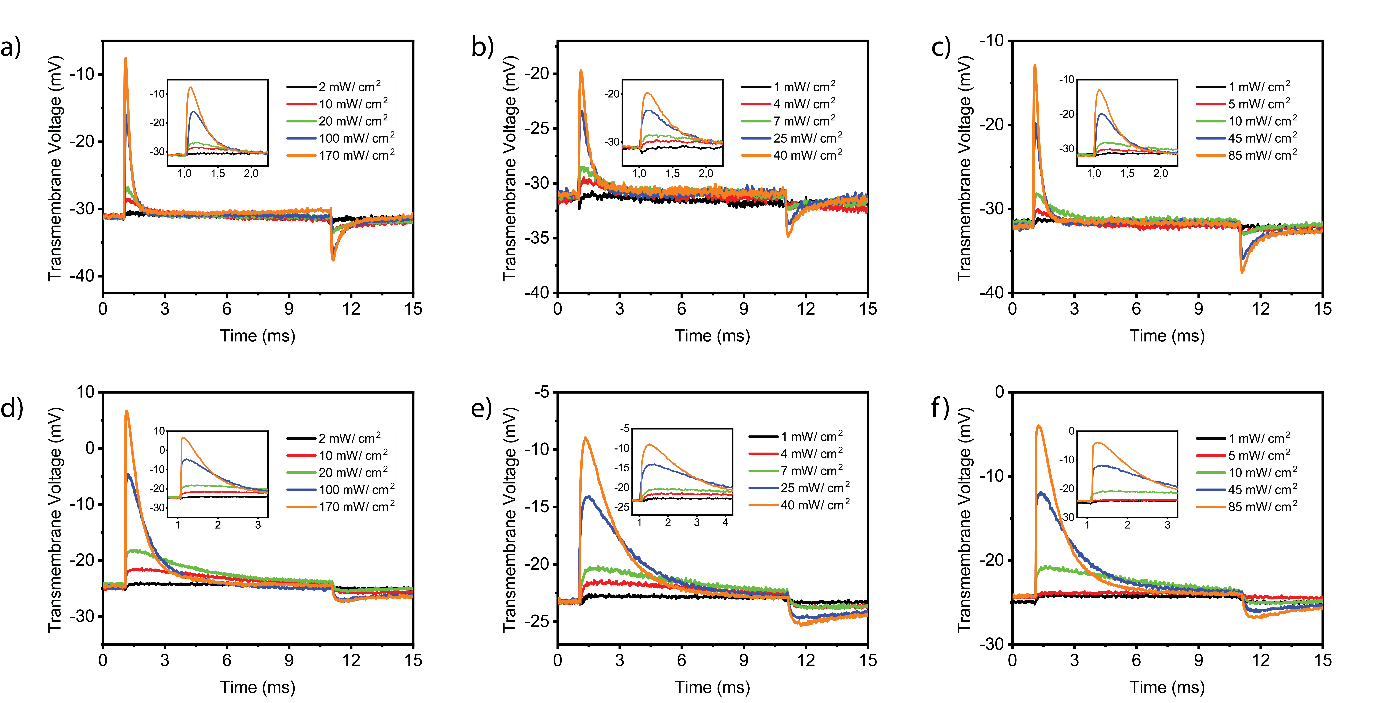

Figure S11. (a)-(c) Transmembrane potential variation of SHSY-5Y cell on control biointerface under light illumination with 10 ms pulse and 100 ms period. (d)-(e) Transmembrane potential variation of SHSY-5Y cell on plasmonic biointerface under light illumination with 10 ms pulse and 100 ms period. ((a,d) – blue light illumination with optical power of 2, 10, 20, 100 and 170 mW. cm^-2^ (b,e) – green light illumination with optical power of 1, 4, 7, 25 and 40 mW. cm^-2^ and (c,f) – red light illumination with optical power of 1, 5, 10, 45 and 85 mW/ cm^-2^ .

Table S7. Depolarization enhancement for both control and plasmonic biointerfaces for blue, green and red light illumination.

|  | Optical power (mW cm^-2^) | Depolarization (mV) | | Enhancement in depolarization (%) | Overall enhancement in depolarization (%) |
| --- | --- | --- | --- | --- | --- |
|  |  | Control | Plasmonic |  |  |
| Blue | 2 | 0.6 | 0.8 | 33.3 | 37.3 |
|  | 10 | 2.5 | 3.6 | 44.0 |  |
|  | 20 | 4.2 | 6.0 | 42.9 |  |
|  | 100 | 15.0 | 19.9 | 32.6 |  |
|  | 170 | 23.4 | 31.3 | 33.8 |  |
| Green | 1 | 0.4 | 0.6 | 50.0 | 25.3 |
|  | 4 | 1.6 | 1.8 | 12.5 |  |
|  | 7 | 2.5 | 3.0 | 20.0 |  |
|  | 25 | 7.6 | 9.0 | 18.4 |  |
|  | 40 | 11.3 | 14.2 | 25.7 |  |
| Red | 1 | 0.5 | 0.6 | 20.0 | 7.7 |
|  | 5 | 1.9 | 2.0 | 5.3 |  |
|  | 10 | 3.3 | 3.5 | 6.1 |  |
|  | 45 | 12.2 | 12.32 | 1.0 |  |
|  | 85 | 19.1 | 20.3 | 6.3 |  |

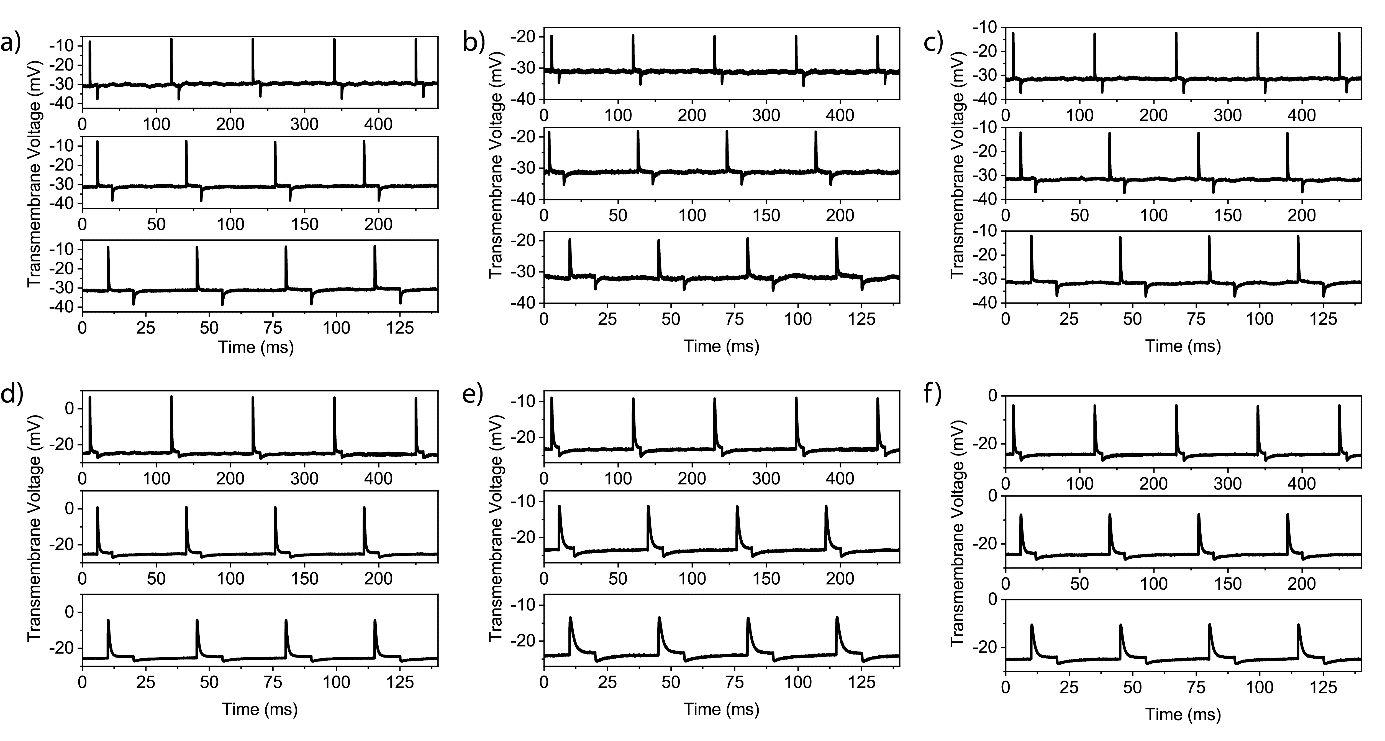

Figure S12. Effect of light evoked pulse trains on control and plasmonic biointerfaces under 10, 20 and 40 Hz illumination (pulse duration 10 ms). (a)-(c) Transmembrane potential variation of SHSY-5Y cell on control biointerface under light illumination with 10ms pulse with 100 ms (10 Hz, top) 50 ms (20 Hz, middle) and 25 ms (40 Hz, bottom) periods. (d)-(f) Transmembrane potential variation of SHSY-5Y cell on plasmonic biointerfaces under LED excitation with 10 ms pulse width with (10 Hz, top), (20 Hz, middle) and (40 Hz, bottom) periods. ((a,d) – blue LED excitation with optical power of 170 mW. cm^-2^ (b,e) – green LED excitation with optical power 40 mW cm^-2^ and (c,f) – red LED excitation with optical power of 85 mW cm^-2^.)

Table S8. Depolarization enhancement for both control and plasmonic biointerfaces for blue, green and red light illumination with 10 ms pulse width and 100 ms, 50 ms and 25 ms periods.

| Light | Frequency (Hz) | Depolarization (mV) | | Enhancement in depolarization (%) | Overall enhancement in depolarization (%) |
| --- | --- | --- | --- | --- | --- |
|  |  | Control | Plasmonic |  |  |
| Blue | 10 | 23.3 | 31.8 | 36.5 | 15.4 |
|  | 20 | 23.4 | 26.7 | 14.1 |  |
|  | 40 | 22.7 | 21.7 | -4.4 |  |
| Green | 10 | 11.2 | 14.5 | 29.5 | 1.3 |
|  | 20 | 12.9 | 12.3 | -4.6 |  |
|  | 40 | 12.7 | 10.5 | -20.9 |  |
| Red | 10 | 19.0 | 20.6 | 10.0 | -9.6 |
|  | 20 | 19.4 | 17.0 | -12.4 |  |
|  | 40 | 19.5 | 14.4 | -26.4 |  |

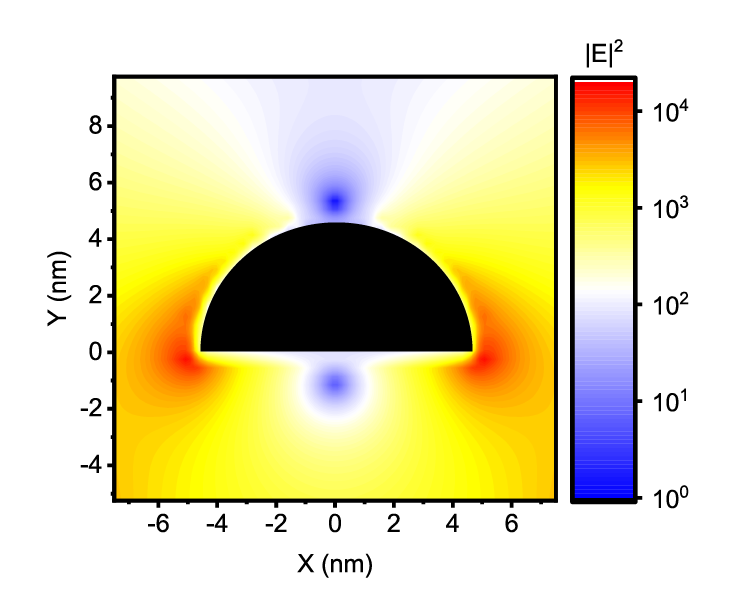

Figure S13. Electric field enhancement profile for AuNI using Lumerical FDTD software.

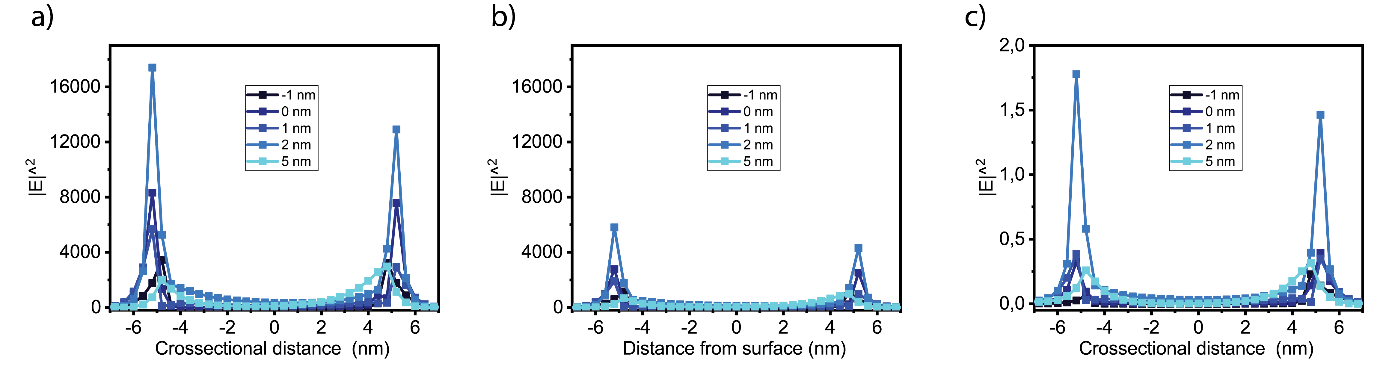

Figure S14. (a) Angle of k vector is 0^0^. The wave illuminated perpendicular to biointerface surface. (b) Angle of k vector is 45^0^. The wave illuminated diagonal to biointerface surface. (c) Angle of k vector is 90^0^. The wave illuminated parallel to biointerface surface.
